# Spin∞ an improved miniaturized spinning bioreactor for the generation of human cerebral organoids from pluripotent stem cells

**DOI:** 10.1101/687095

**Authors:** Alejandra I. Romero-Morales, Brian J. O’Grady, Kylie M. Balotin, Leon M. Bellan, Ethan S. Lippmann, Vivian Gama

## Abstract

Three-dimensional (3D) brain organoids derived from human pluripotent stem cells (hPSCs), including human embryonic stem cells (hESCs) and induced pluripotent stem cells (iPSCs), have become a powerful system to study early development events and to model human disease. Cerebral organoids are generally produced in static culture or in a culture vessel with active mixing, and the two most widely used systems for mixing are a large spinning flask and a miniaturized multi-well spinning bioreactor (also known as Spin Omega (SpinΩ)). The SpinΩ provides a system that is amenable to drug testing, has increased throughput and reproducibility, and utilizes less culture media. However, technical limitations of this system include poor stability of select components and an elevated risk of contamination due to the inability to sterilize the device preassembled. Here, we report a new design of the miniaturized bioreactor system, which we term Spin∞ that overcomes these concerns to permit long-term experiments.

## Introduction

Brain organoids are three-dimensional (3D) structures formed from neural stem cells (NSCs) derived from human pluripotent stem cells (hPSCs) that can effectively model human brain development up to 12-14 weeks post-conception [1–4], a time period which includes critical patterning events in the cerebral cortex and other brain regions [5,6]. On a cellular level, brain organoids show a high level of similarity to the *in vivo* developing human brain in the early stages of development, including progenitor zones (ventricular zone and subventricular zone consisting of PAX6+/SOX2+ NSCs) that form around central lumens [1,4]. These 3D organoid cultures therefore provide a robust system amenable to extended cultivation and manipulation, which makes them a useful tool to model development and disease in the context of the complex brain microenvironment [7–9].

Recently, two protocols have been published on enhancing cortical plate formation within hPSC-derived cerebral organoids: one using large spinner flasks and microfilaments as a solid support [1,10] and another that uses a miniature spinning bioreactor termed Spin Omega (SpinΩ) [11,12], which consists of 3D printed gears and paddles driven by a single electric motor. The SpinΩ provide an accessible and versatile format for culturing brain-region-specific organoids due to its reduced incubator footprint, decreased media consumption, and increased throughput, but several technical caveats limit its use in long-term experiments, most prominently the choice of components used to fabricate the device and the design of the device with respect to limiting the chances of contamination and mechanical failure. Here, we redesigned the SpinΩ to overcome these problems, leading to the creation of a device that we have termed Spinfinity (Spin∞).

## Results and Discussion

This improved spinning bioreactor is customized for organoid culture via miniaturization and increased throughput. The device has a small footprint and an integrated spinning mechanism that does not require a large amount of dedicated incubator space (Figures 1-16). Organoids can be grown under many different conditions in parallel, with each condition requiring a minimal culture medium to reduce cost. In addition, the device is biocompatible and sterilizable for repeated use. To develop this improved device, we first considered the choice of motor that drives the spinning bioreactor. Motor life span inside an incubator can be a major hurdle when using a bioreactor system because motors that are not designed to withstand harsh environments (high humidity, 5% carbon dioxide, and elevated heat) can easily corrode and break, leading to unexpected failures in the middle of long-term experiments. We therefore selected a motor with the ability to operate at increased temperatures (max 70°C) and humidity (90% humidity). Additionally, parylene vapor deposition was applied to the motor to provide an additional moisture barrier to further the lifetime of the motor and increase durability [13]. Next, we considered the basic design of the bioreactor. Due to the humidity issue raised above, we used stainless steel screws, standoffs, nuts, and washers in order to reduce the oxidation of the metal and prolong the life of the bioreactor. These parts also have the advantage of being autoclavable as separate components or with the assembled bioreactor, and the updated design now allows the majority of the equipment (including the 3D printed parts) to be assembled and autoclaved to reduce external handling and improve sterility. The addition of an upper acrylic lid on the bioreactor (where the motor is anchored) further enhances stability and consistency of the device. The SpinΩ motor was anchored in two points on the upper lid, which frequently caused the motor to shift and oscillate, thereby putting unnecessary additional stress on the motor leading to eventual failure. Because the updated Spin∞ was designed using a separate acrylic sheet, hex standoffs, and a larger, more durable motor, all components are kept in perpendicular alignment to the well plate, thus preventing shifting and oscillation. Additionally, because the Spin∞ is designed with hex standoffs and an autoclavable 3D printed base, the lid is securely placed on the 12-well plate, which prevents spills and possible contamination points. By comparison, the original SpinΩ design freely rests on the top of the plate, making this design prone to spillage and contamination. Overall, these changes yield a miniaturized spinning bioreactor that is exceptionally easy to maintain and performs consistently. This updated device is amenable to months-long experiments without concern of unexpected malfunctions.

**Figure 1.**
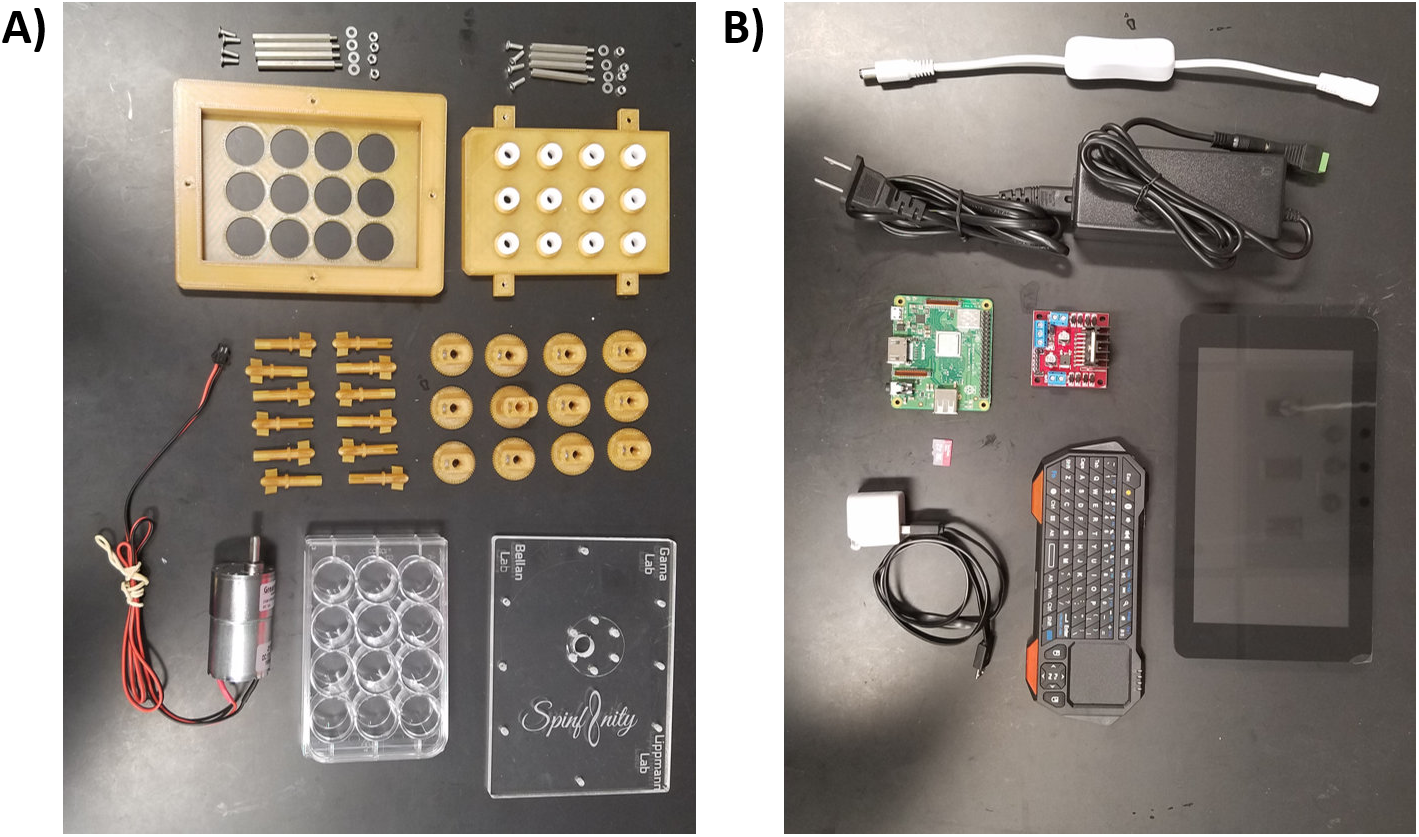
All individual components used to build the Spin∞. **A)** Hardware and 3D printed components. **B)** Electronic components.

**Figure 2.**
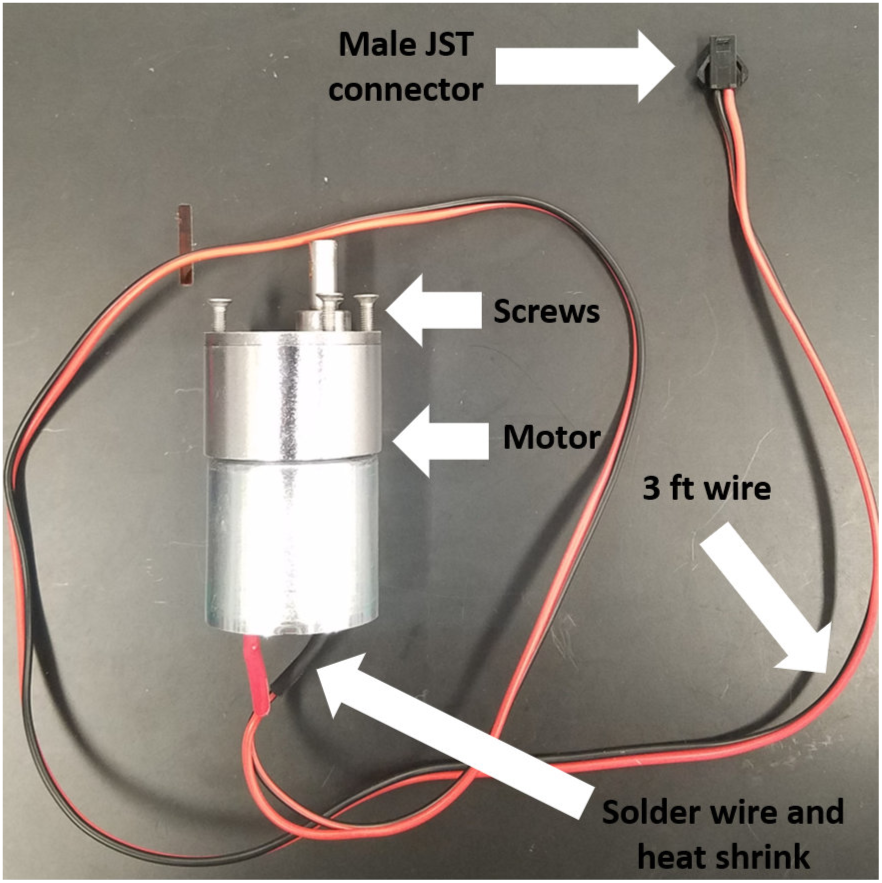
Motor preparation.

**Figure 3.**
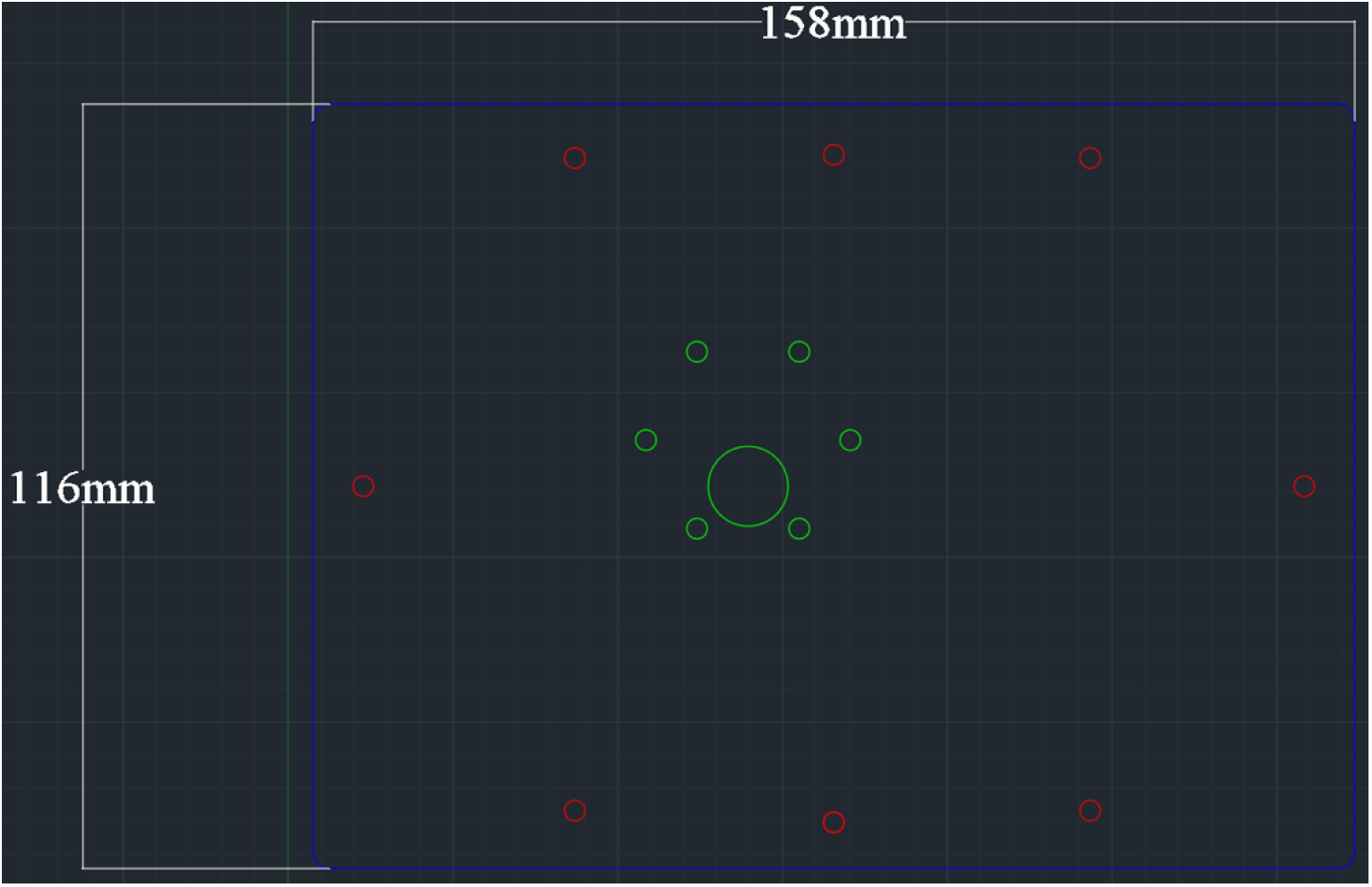
Acrylic plate dimensions for laser cutting.

**Figure 4.**
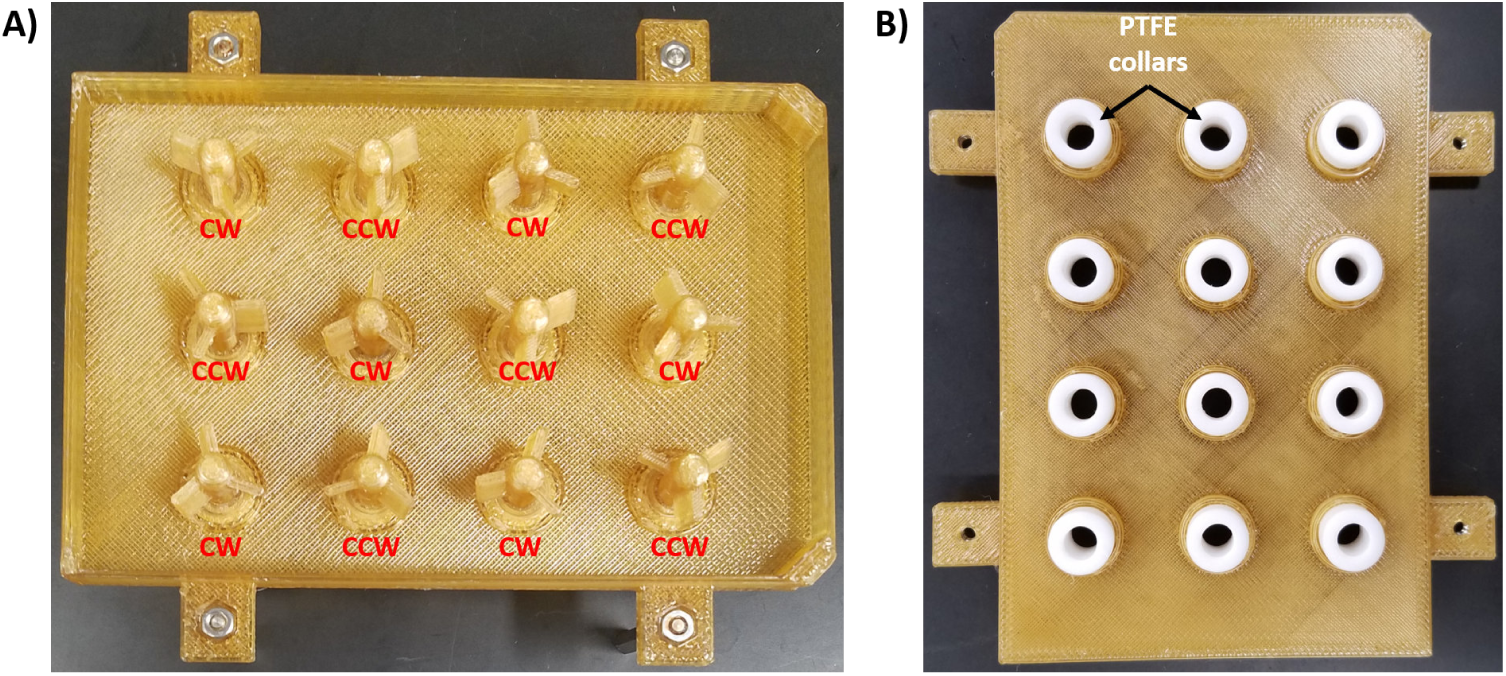
Insertion of paddles into the 12-Well Plate Lid. **A)** Positioning of the CW Paddles and CCW Paddles are noted. **B)** Positioning of the PTFE collars are shown on the opposite side of the lid.

**Figure 5.**
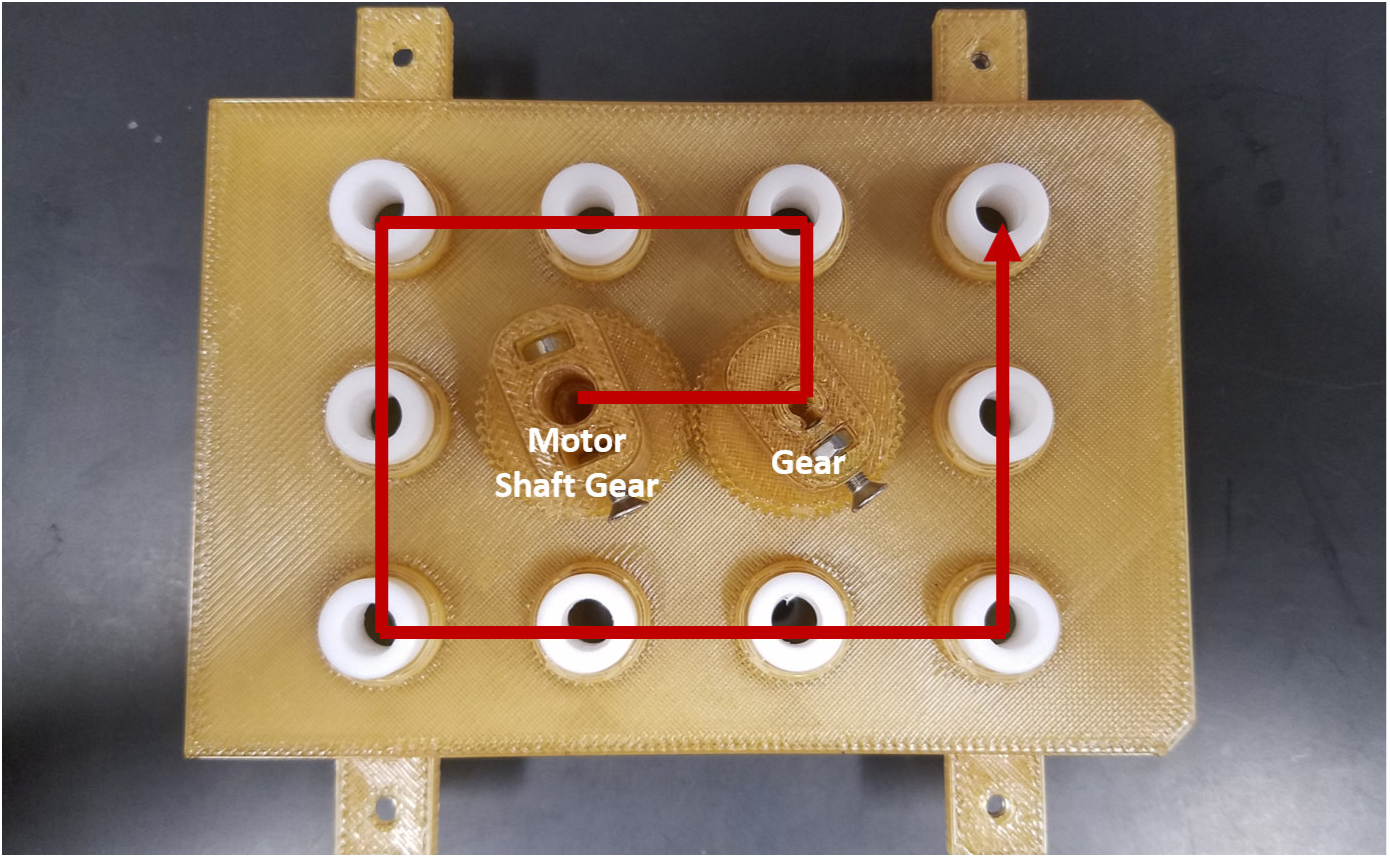
Assembly pattern of the Gears to ensure proper operation. Make sure each gear is properly threaded before moving on to the next one. The position of the Motor Shaft Gear is noted.

**Figure 6.**
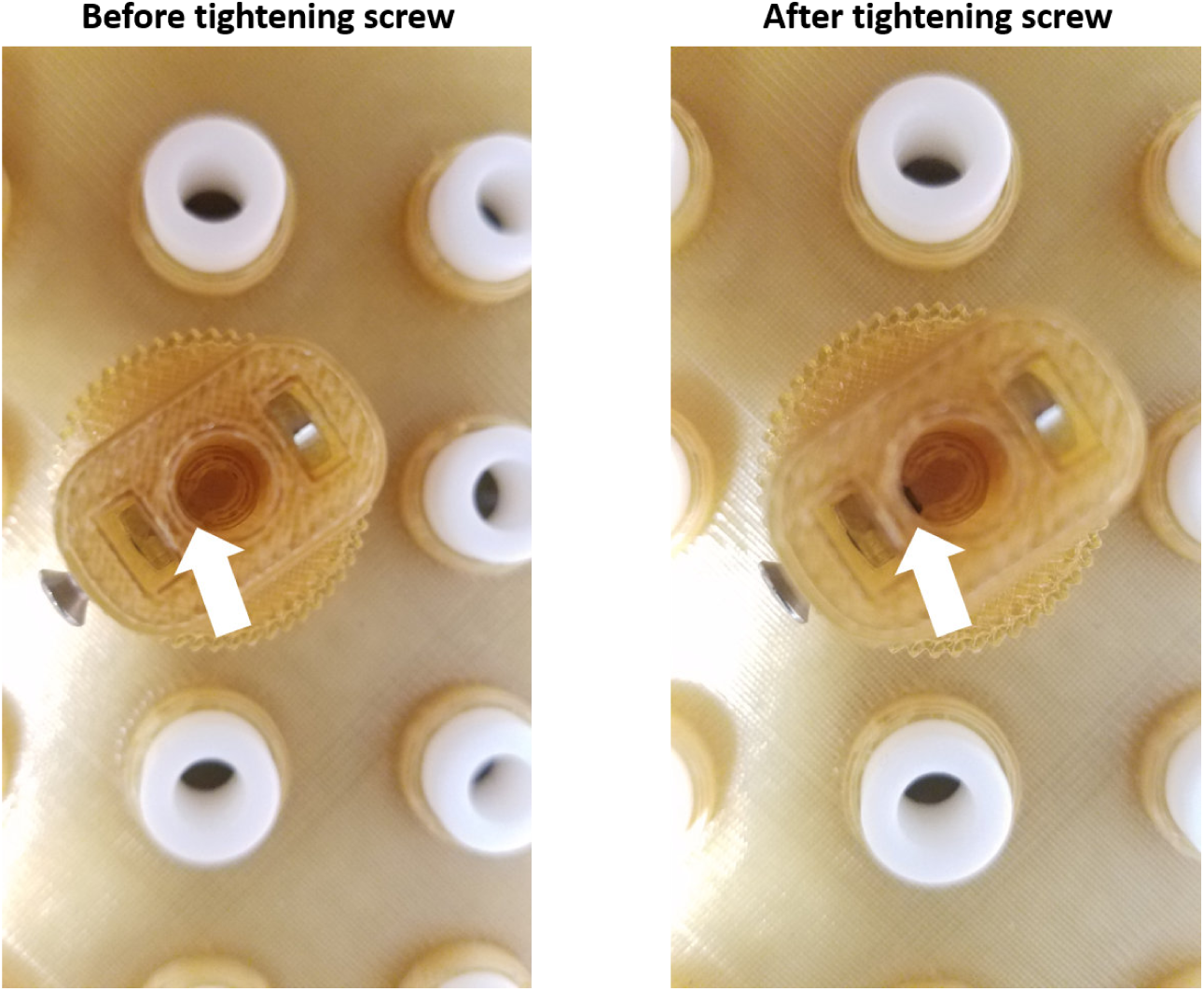
Proper positioning of screws in the Gears. Screws should not be overtightened.

**Figure 7.**
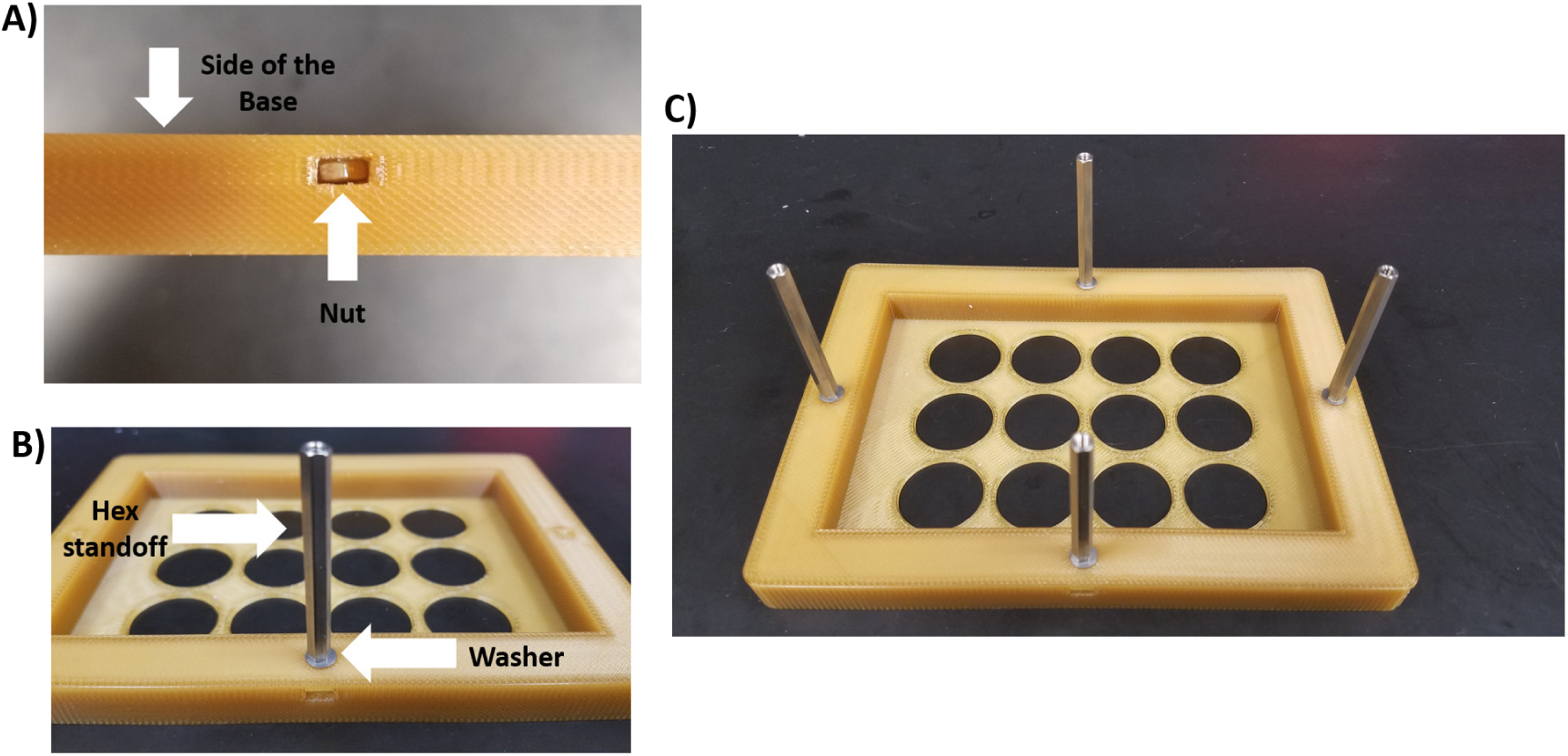
Attachment of the 45 mm hex standoffs to the Base. **A)** Each nut is placed into the side of the Base. **B)** A washer is added to the 45 mm hex standoff, which is then screwed into the nut. **C)** Image showing all hex standoffs attached to the Base.

**Figure 8.**
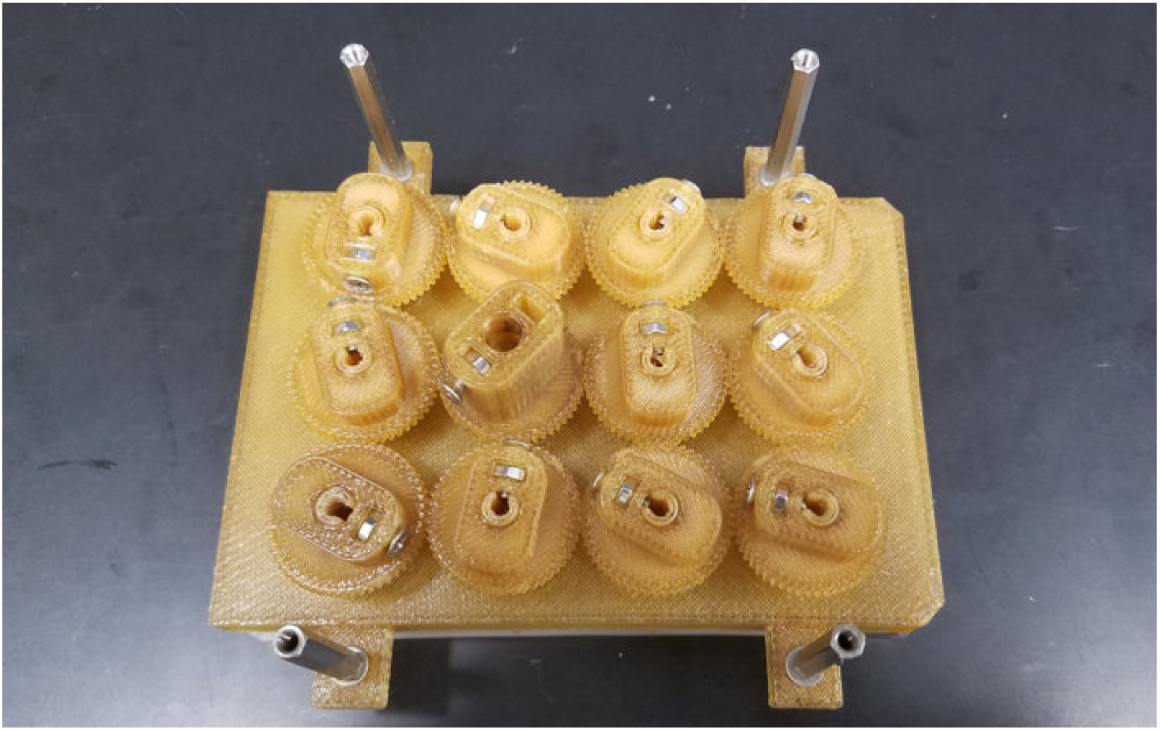
Attachment of the 35 mm hex standoffs to the 12-Well Plate Lid.

**Figure 9.**
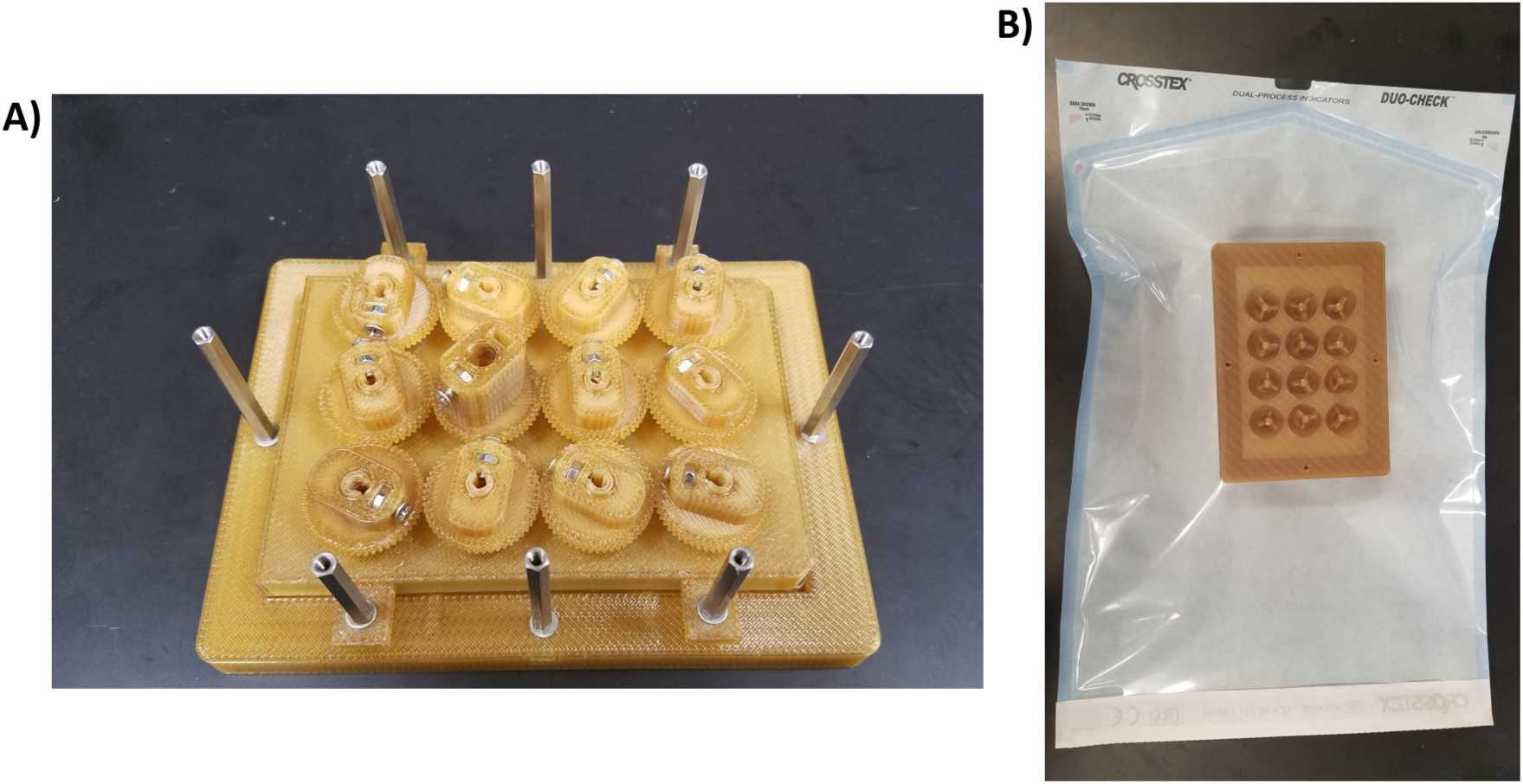
Preparation of the bioreactor for autoclaving. **A)** The Base with the hex standoffs, the 12-Well Plate Lid with the Paddles and Gears, and the stainless-steel screws can all be autoclaved. **B)** The piece should be placed upside down in an autoclavable bag (the Base should face the clear side of the bag).

**Figure 10.**
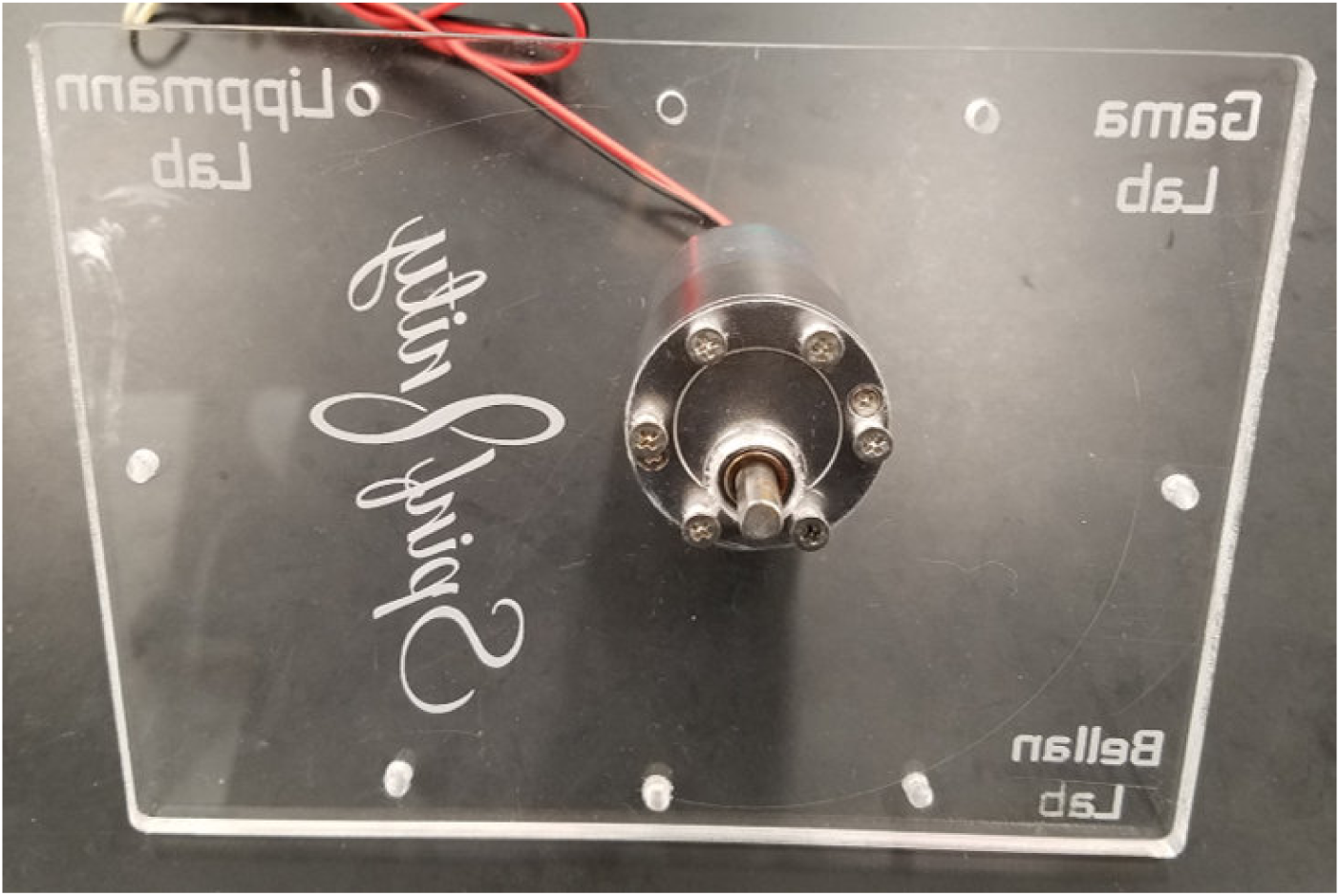
Attachment of the motor to the acrylic plate.

**Figure 11.**
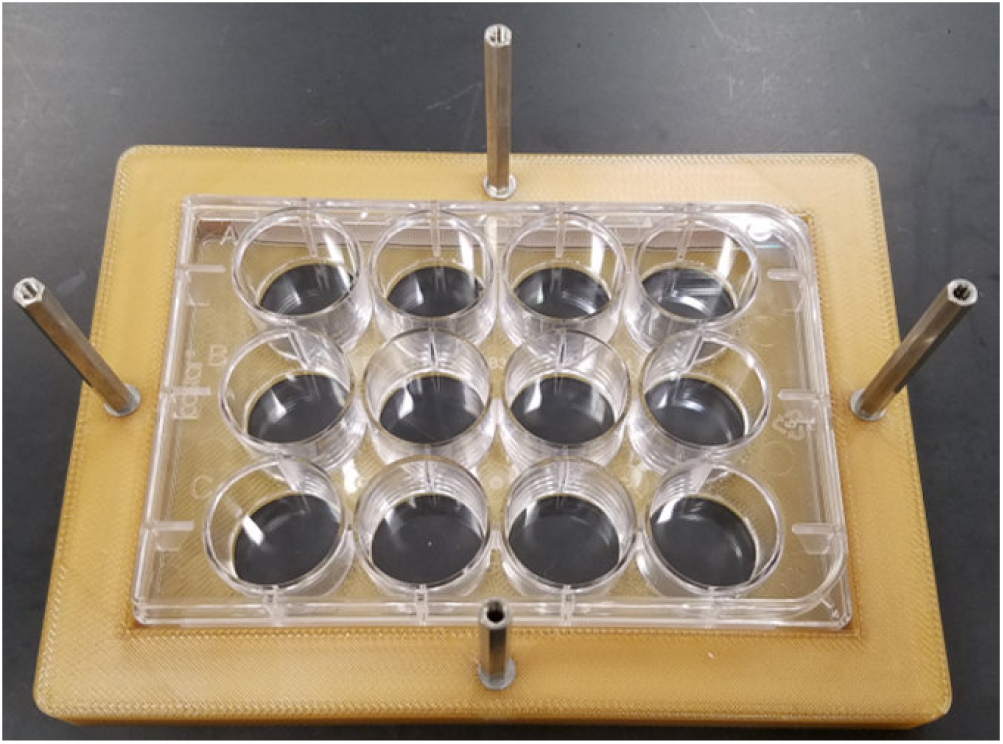
Placement of a 12-well plate into the Base.

**Figure 12.**
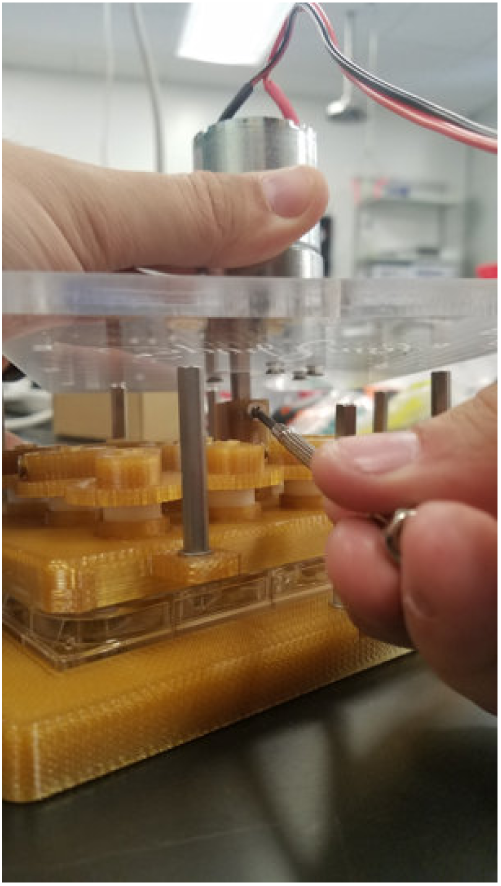
Attachment of the beveled side of the motor shaft into the Motor Shaft Gear.

**Figure 13.**
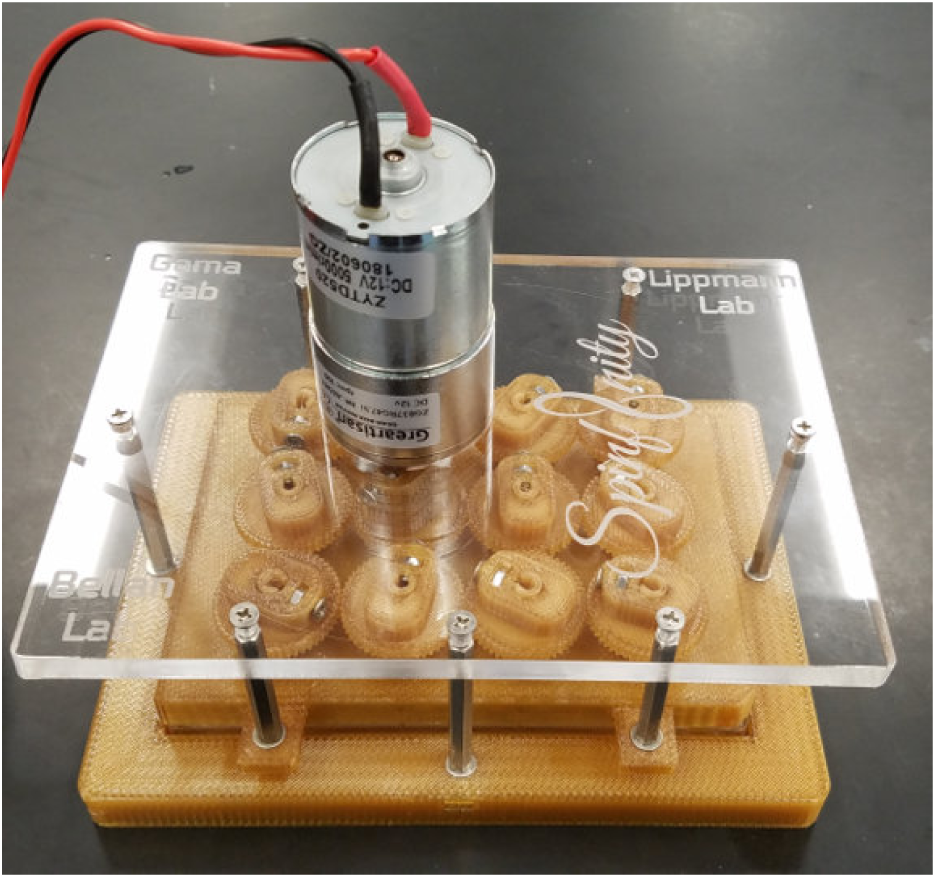
Fully assembled bioreactor.

**Figure 14.**
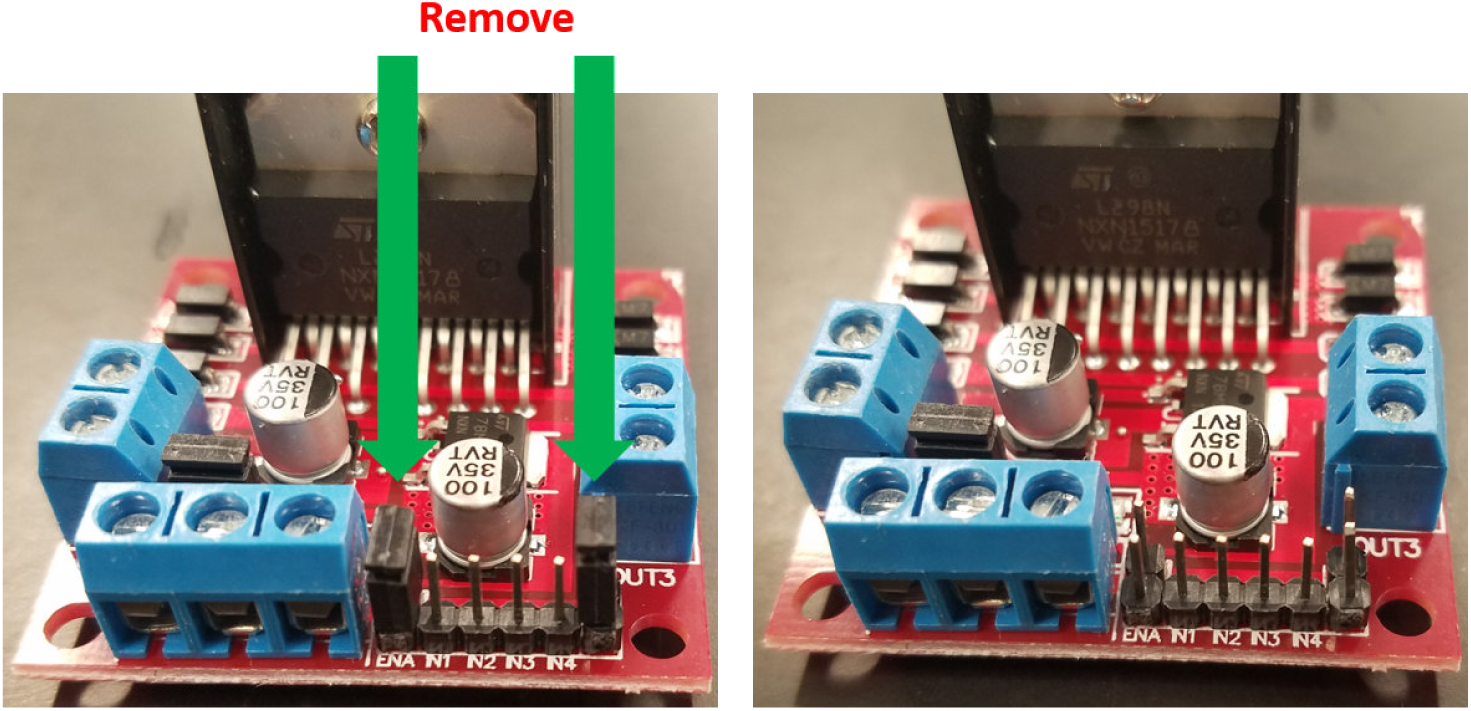
Removal of the jumpers from the L298n bridge.

**Figure 15.**
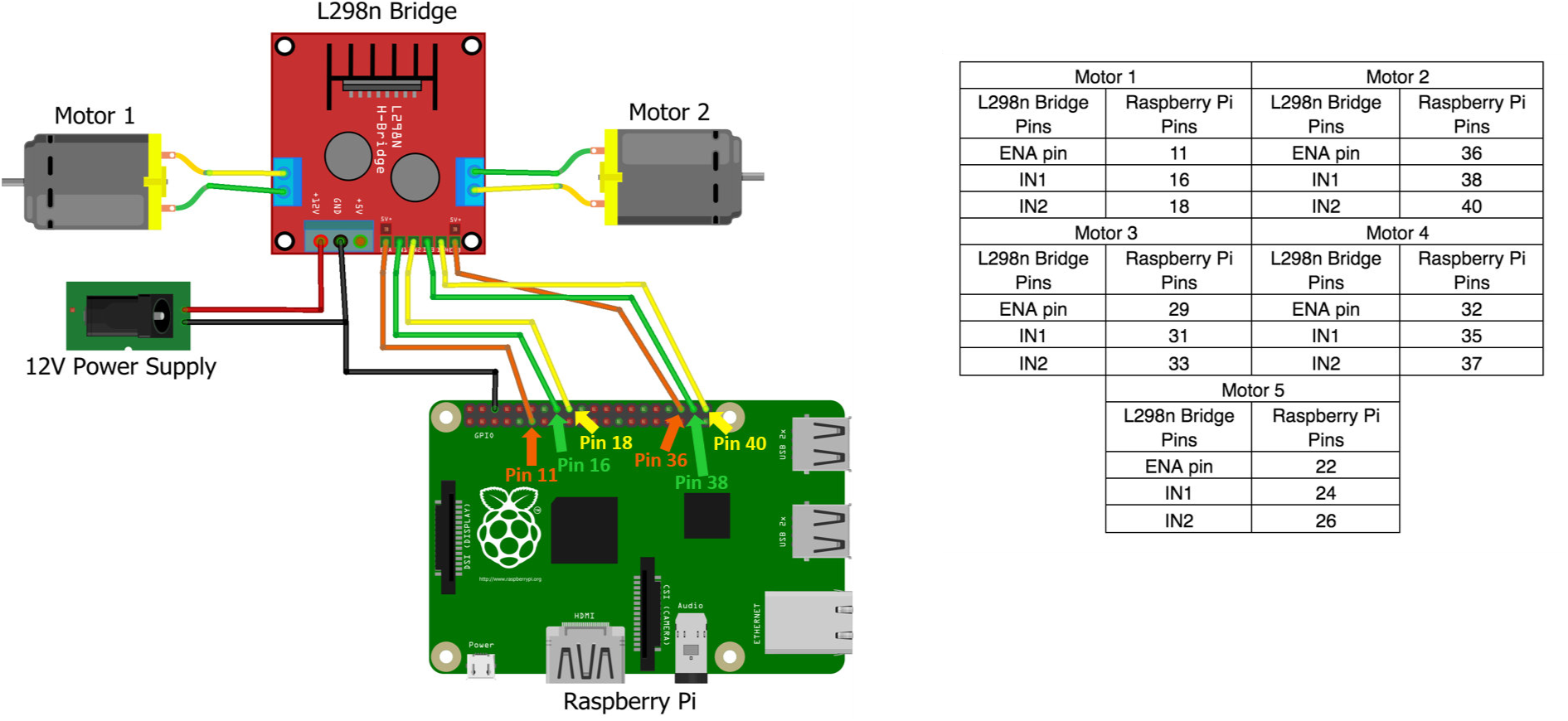
Example for how to connect an L298n bridge to two motors and a Raspberry Pi. The accompanying table provides pin locations for connecting the Raspberry Pi to up to five motors.

**Figure 16.**
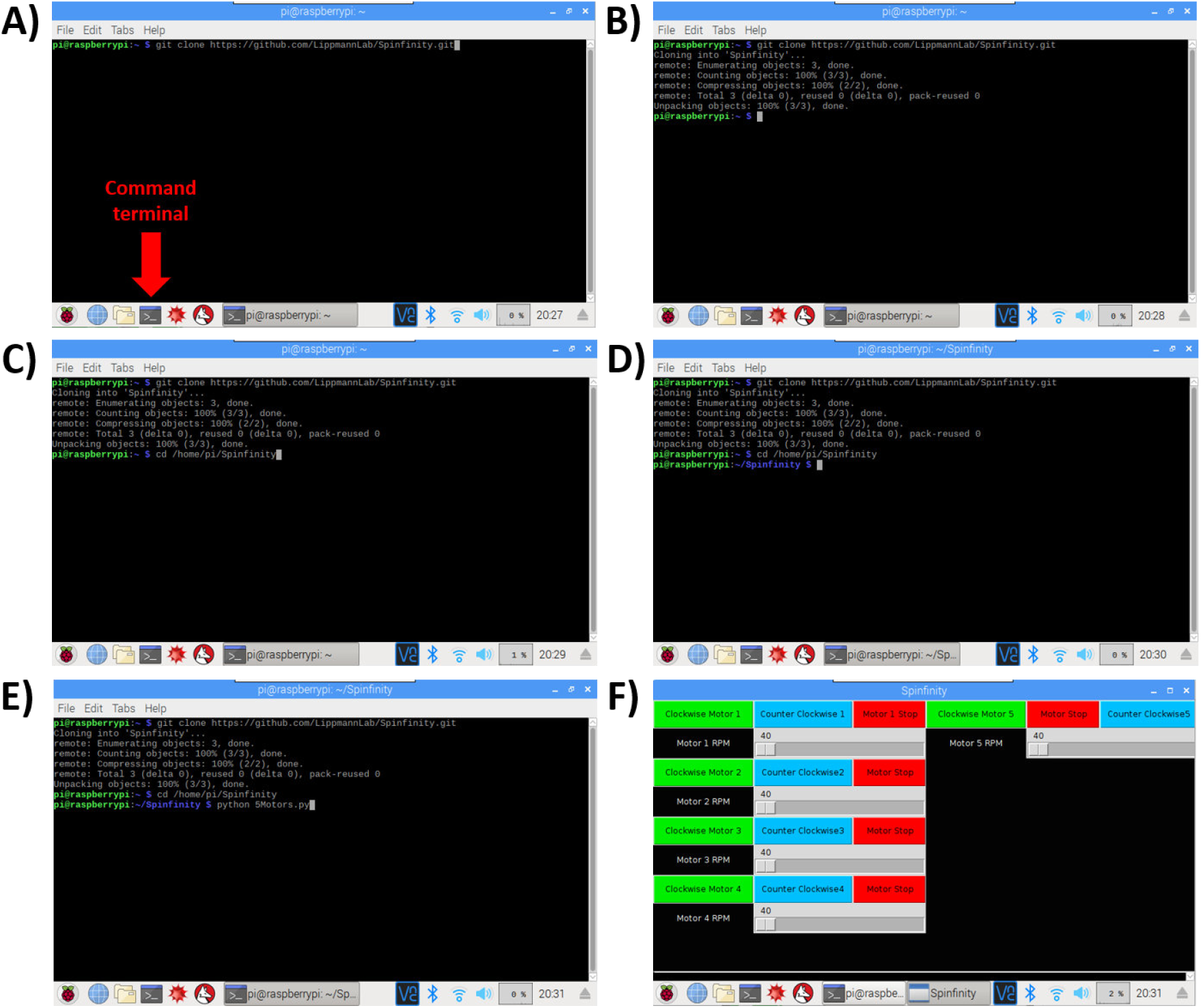
Software installation setup. **A-B)** The Command Terminal is opened, and by entering the text shown in the figure, the repository will be cloned from Github into a folder on the Raspberry Pi. **C-D)** Opening of the repository. **E-F)** Opening of the software, which leads to the user interface with touchscreen control.

Brain organoids were generated using the STEMdiff™ Cerebral Organoid Kit (Stem Cell Technologies) with some modifications (Figure 17A and 17B) [1,8,11,13]. As the generation of homogeneous embryoid bodies (EBs) is critical for achieving homogenous organoids, we utilize a 24-well plate AggreWell™ 800 (Stem Cell Technologies, catalog 34815) [14,15] to increase the reproducibility and the yield of EBs (Figure 18A and 18B). After 4 days of culture in the AggreWell, we transfer the organoids to a 10 cm ultra-low attachment plate to allow further growth of the EBs before the Matrigel embedding process (Figure 17B). The diameter of the EBs increased over time; by the embedding day (day 7) the average diameter was 387 μm (±57) and by the bioreactor transfer day (day 10) the average diameter was 661 μm (±112) (Figure 18B). High resolution images show the formation of organized neural rosettes within the EBs at day 8 [1,11]. Alpha-tubulin (α-tubulin filaments) can be seen organized radially from the lumen (Figure 18C) [16–19]. These polarized neuroepithelium-like structures resemble neural tube formation and are the precursors to the formation of the brain lobules [10].

**Figure 17.**
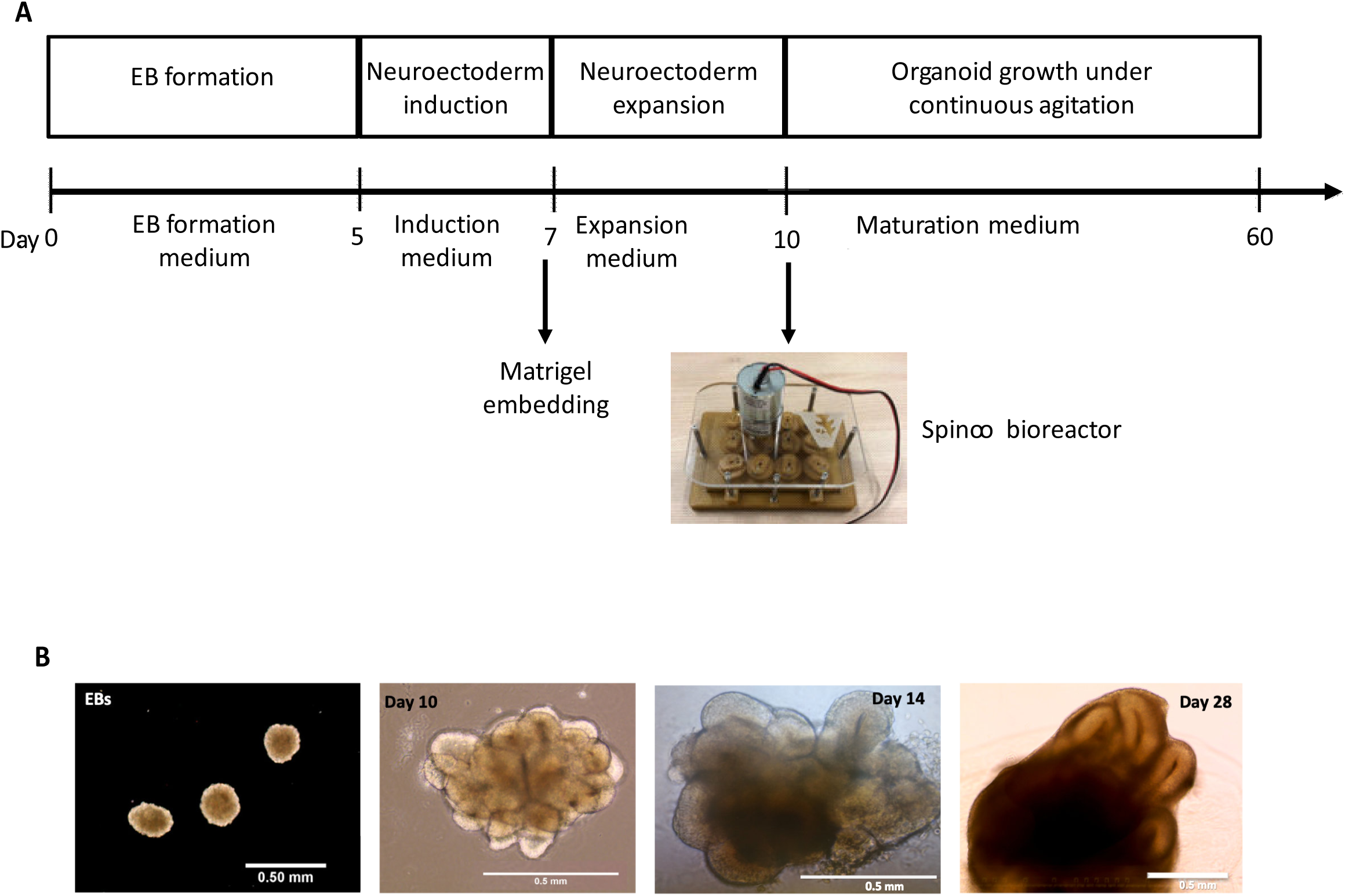
Protocol for the generation of brain organoids using Spin∞. **A)** Schematic of the major stages in the culture protocol. **B)** Macroscopic images of the organoid growth and development. Scale bars: 0.5mm.

**Figure 18.**
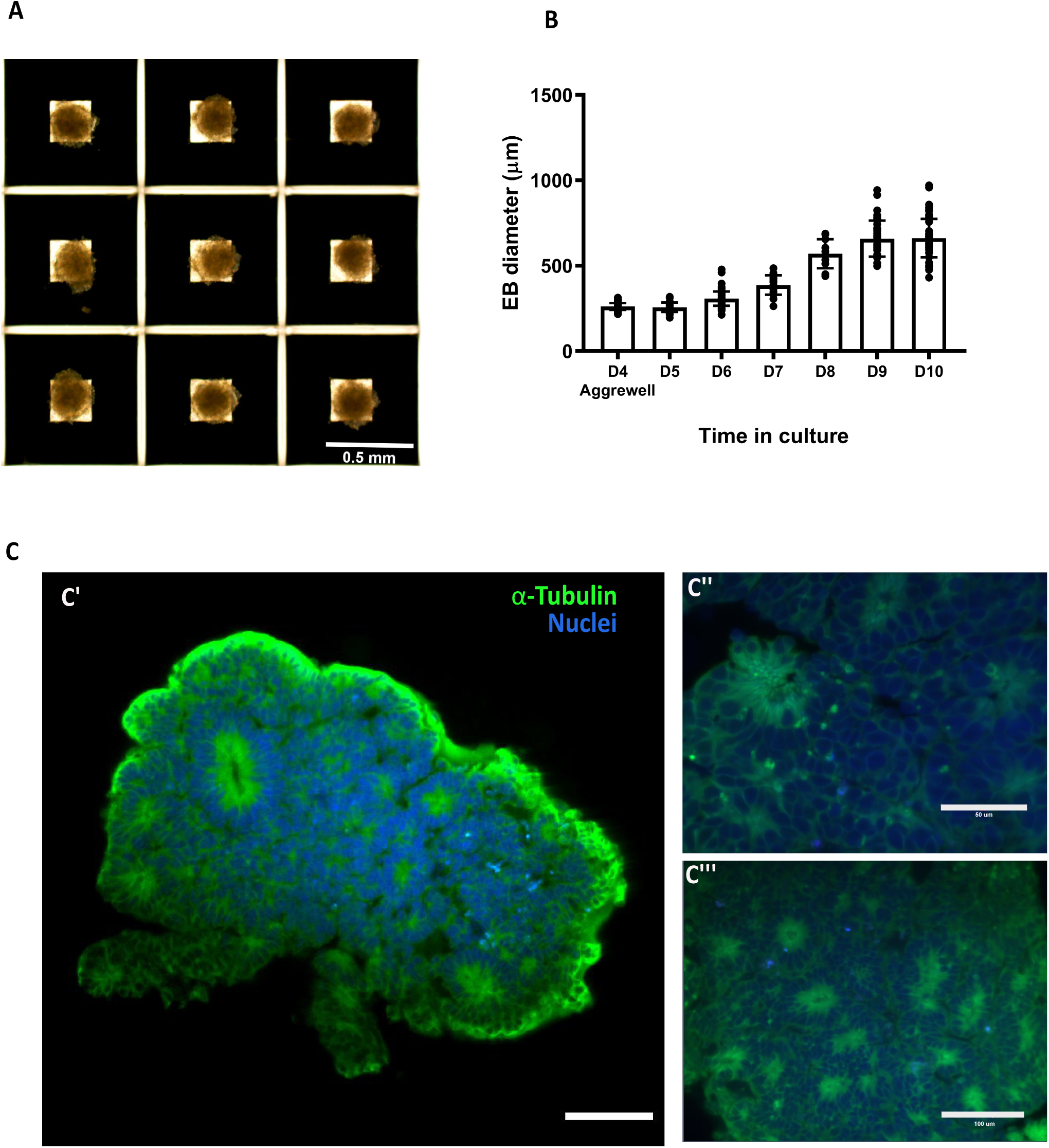
Embryoid body generation and size over time. **A)** Four days old Embryoid bodies generated using microwells. **B)** Relative growth of the embryoid bodies over time. **C)** Light sheet microscope images of day 7 embryoid body showing the formation of organized neural rosettes. C’’ and C’’ show close up to the rosettes Data are mean ± s.d. Scale bars: (A) 0.5mm, (C’) 100 μm, (C’’) 50μm, (C’’’) 100μm.

Characterization of the organoids was performed at day 60 and day 150. Organoids were fixed in 4% PFA for 15 min at 4°C, followed three 5 min washes in PBS and 30% sucrose dehydration overnight at 4°C. Embedding for sectioning was performed as reported previously [1]. Cryosections (15 μm thick) were stained using neural progenitor cell (NPC), mature pan-neuronal, and cortical layer markers. As previously reported for day 60 organoids [11], the cells that stain positive for neural progenitor markers are localized in the periphery to the ventricle-like structures (Figure 19A). Multi-layer stratified structures can be readily seen and are comprised of NPC+ cells marked by SOX1, SOX2, and PAX6 marking the progenitor zone (Figure 19A-19C). PAX6 expression confirmed the forebrain identity in the regions of interest [1]. Nestin positive cells mark radial glia, key structures for the expansion of the mammalian cortex by differentiating into neurons and intermediate progenitor cells (Figure 19A and 19B) [20].

**Figure 19.**
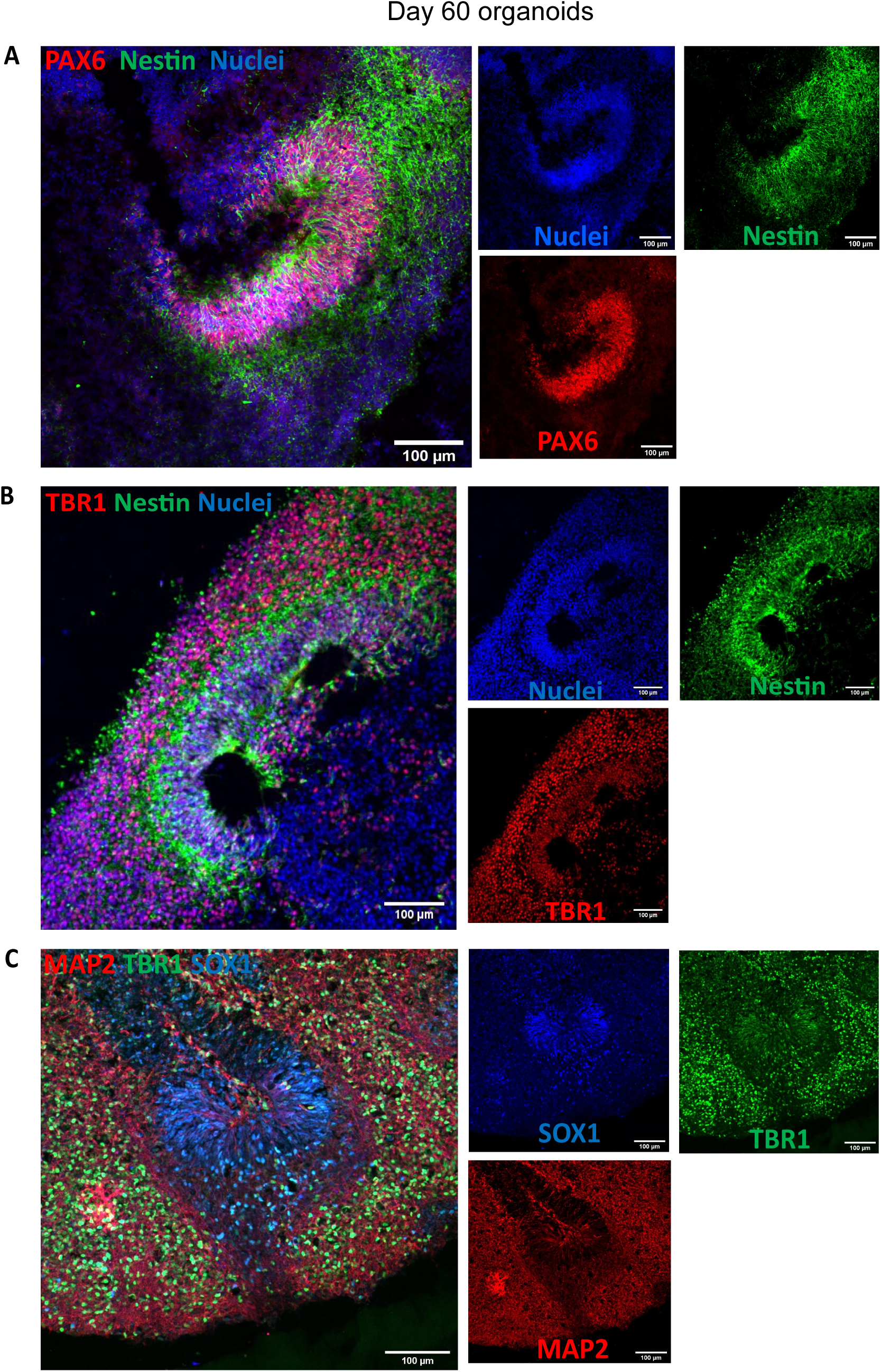
Staining for neural progenitor cells and cortical neurons. **A)** Day 60 organoids showing the presence of neural progenitor markers Pax6 and Nestin. **B)** Cortical plate marker TBR1 shows the organization of the pre-plate. **C)** MAP2 positive cells indicate the presence of committed neurons at this stage. Scale bars: (A-C) 100μm

Pre-plate formation was confirmed by the presence of TBR1+ cells (Figure 19B and 19C). This marker also identifies cells localized to the early-born layer VI of the cortex [21]. TBR1+ neurons are vital for guiding the subsequent neuronal migrations [21]. Radial organization can be seen by MAP2, a neuronal marker for dendritic outgrowth and branching (Figure 19C) [22]. Cajal-Retzius cells, a cell population crucial to the generation of the cortical plate architecture, is present by Reelin+ neurons located along the organoid surface (Figure 20A). These early born cells localize to the marginal zone of the cortex (layer I) and contribute to the formation of the inside-out layering of the neurons in the neocortex [23].

**Figure 20.**
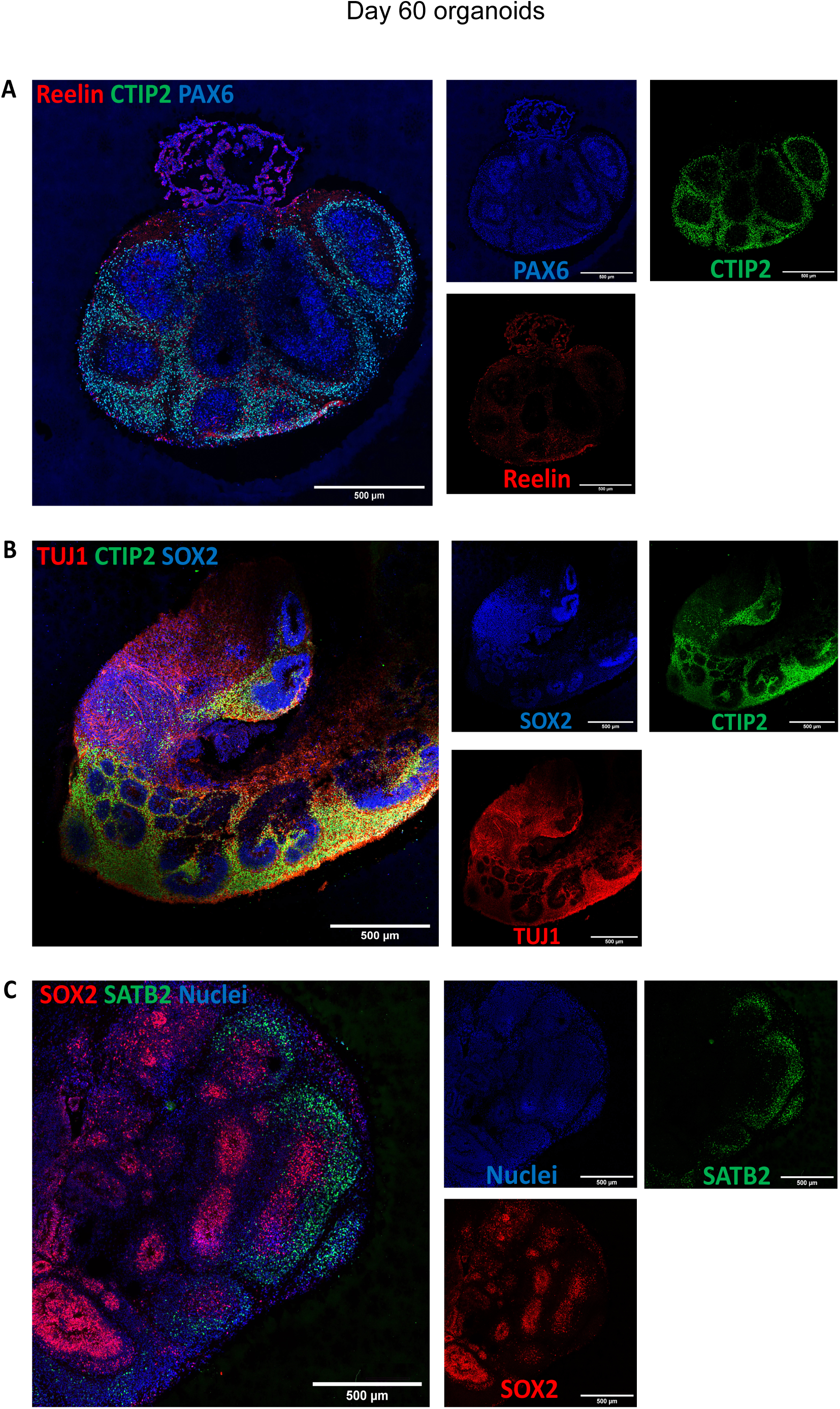
Characterization of cortical layers present after 60 days in culture. **A)** Cajal-Retzius neurons stained for reelin show the presence of marginal zone. **B)** Staining for deep layer neurons (CTIP2) and neuronal markers (TUJ1). **C)** Upper-layer marker SATB2 indicate the presence of neurons belonging to the cortical layer IV. Stitched images at 20X, scale bars: 500μm

At day 60, early-born neurons from layer V were positive for CTIP2 (also known as BCL11B) (Figure 20A and 20B). Furthermore, late-born neurons from the superficial layers (layer IV) can be seen as SATB2+ cells (Figure 20C). Interestingly, these markers show a clear separation from the neural progenitor zone, indicating a spatial separation of the different neuronal lineages, as well as the recapitulation of the cortical architecture observed in other brain organoid protocols.

Neurons expressing layer II/III markers CUX1 and BRN2 are present by day 150 (Figure 21A and 21B). Presence of this late born neurons [1,5,11,24] underlie the capacity of the organoid system to recapitulate the cytoarchitecture of the developing cortex.

**Figure 21.**
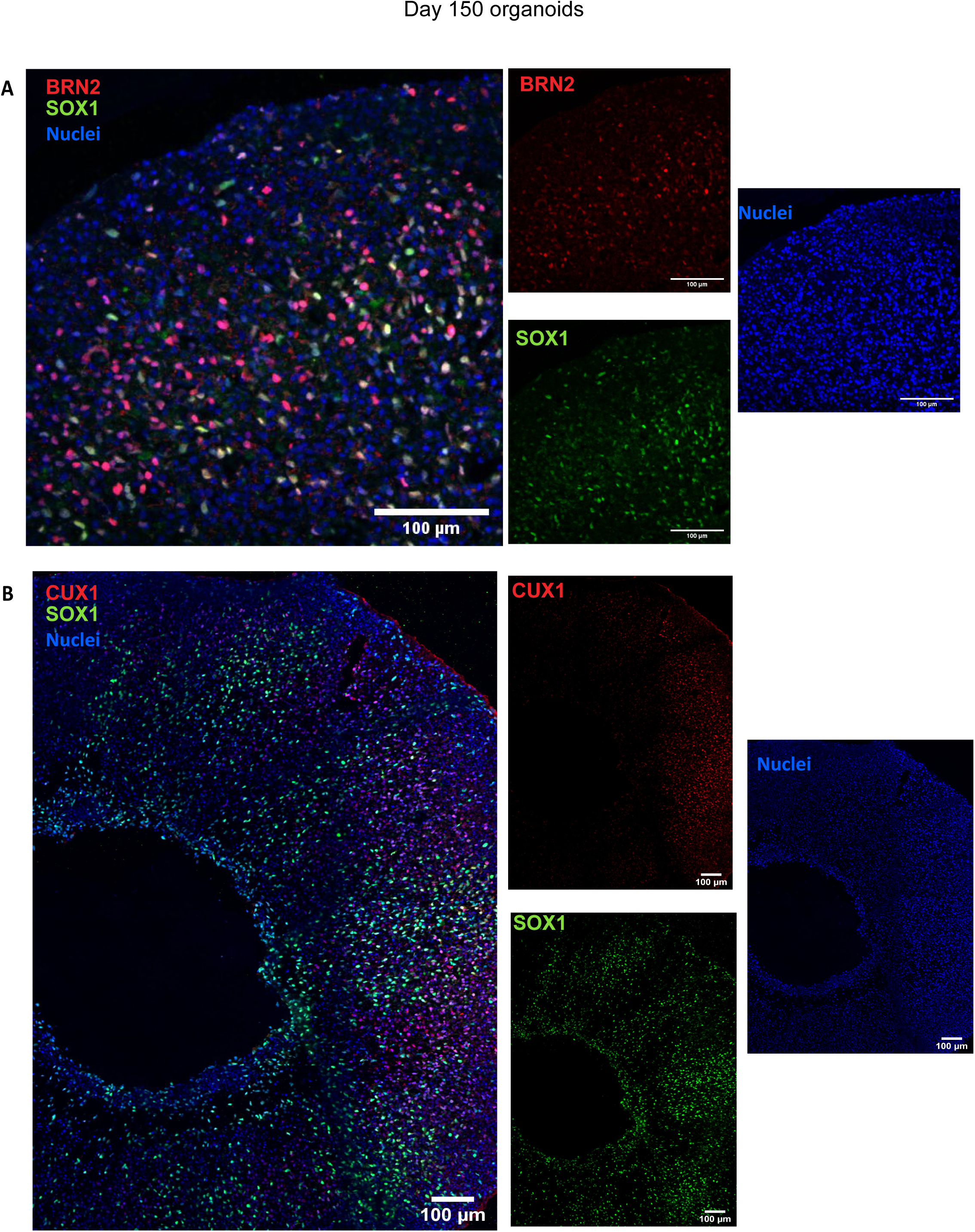
Long term culture of brain organoids allows upper layer specification. Sample images of immunostaining for superficial layer neuron markers **(A)** BRN2 and **(B)** CUX1 in cerebral organoids at day 150. Scale bars: 100 μm.

Finally, we analyzed the expression of the apoptosis marker cleaved Caspase 3 (ClC3). At day 60 and day 150, the presence of apoptosis is evident but the integrity of the cortical structures is maintained (Figure 22A and 22B). Apoptosis is a key process during brain formation, controlling cellularity in the developing brain [25,26]. Work in human [27], mouse [26] and rat [28] brain samples shows high incidence of cell death during the development of the neocortex. As the overall architecture of the organoids was maintained, the observed cell death may be the result of the normal elimination of cells that takes place in the developing brain.

**Figure 22.**
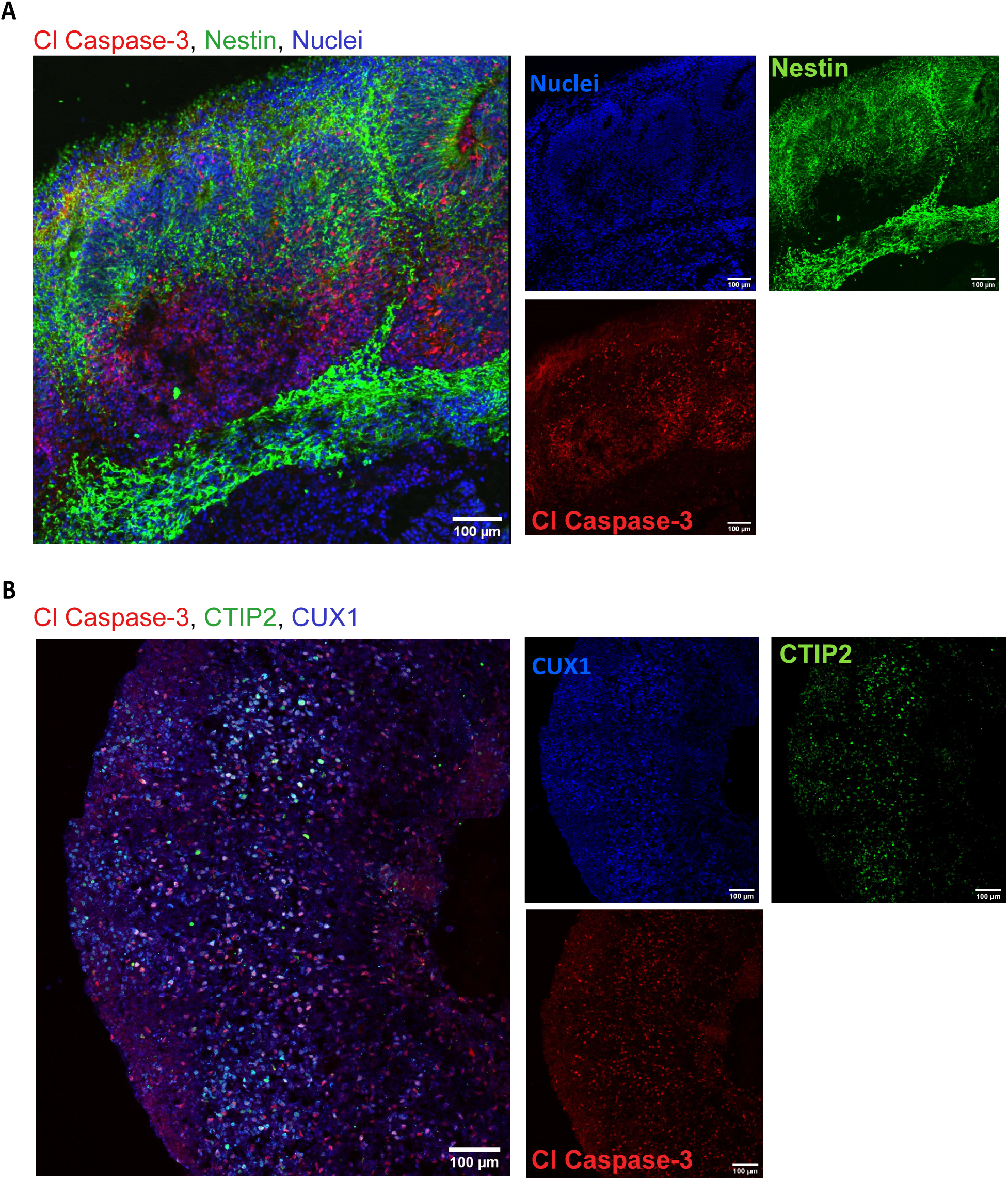
Immunostaining for apoptosis marker Caspase 3. **A)** Cells positive for cleavage caspase 3 at day 60 **B)** and day 150 show no structural compromise of the cortical architecture. Scale bars: 100μm. (B) stitched image.

Here we have shown that organoids cultured Spin∞ have the growth capacity and laminar organization previously reported in the literature [1,8,11,13]. Brain organoids grow above 3 mm in diameter and have distinct and organized segregation between all cortical layers. Human NPC markers, as well as deep cortical and pan-neuronal markers can be identified in a structured manner and in the expected stages. Evaluation of the apoptotic marker ClC3 shows cell death in the core of the organoid as expected with no major compromise of the organoid integrity.

While the data shown here only include brain organoids grown up to day 150, we have successfully culture them over 200 days without the need to change motors, which demonstrates the increased motor life span of Spin∞ under high temperature and humidity conditions. In addition, no contamination was detected even after long-term culture, which we suggest is due in part to the ability to autoclave the majority of Spin∞ components after assembly. Although we find the protocols described here to be highly reproducible, they do not generate completely uniform organoids but rather contain a cluster of individual protruding cortical units. Similar observation was reported with the original SpinΩ.

## Methods

### Human pluripotent stem cells

We have successfully used feeder-dependent hiPSCs reprogrammed from newborn human foreskin fibroblasts (ATCC, cat. no. CRL-2522), as well as human embryonic stem cells (H9 from WiCell). Human iPSC lines AICS-0012 (WTC-mEGFP-TUBA1B-cl105) used in the current study were obtained from the Allen Institute (Coriell Institute for Medical Research, New Jersey). This cell line has a N-term insertion of mEGFP in frame generated by CRISPR-Cas9 technology. Induced pluripotent stem cells were grown in feeder-free conditions in plates coated with Matrigel (Corning) and maintained in E8 media at 37°C with 5% CO2. Culture medium was changed daily. Cells were checked daily for differentiation and were passaged every 3-4 days using Gentle dissociation solution (Stem Cell Technologies). All experiments were performed under the supervision of the Vanderbilt Institutional Human Pluripotent Cell Research Oversight (VIHPCRO) Committee (Record ID#56).

### Brain organoids

Brain organoids were generated using the STEMdiff™ Cerebral Organoid Kit (Stem Cell Technologies) using the manufacturer protocol with some modifications [1,8,11,13]. On Day 0, iPSCs were detached with Gentle Cell Dissociation Reagent for 8 minutes at 37°C. Cells were resuspended in 1mL of EB Seeding Medium and centrifuged at 300g for 5 minutes. The cell pellet was resuspended in 2mL of EB Seeding Medium. Homogeneous and reproducible EBs were generated by using 24-well plate AggreWell™ 800 (Stem Cell Technologies, catalog 34815). In each well, 2.7×106 cells were plated (approx. 9000 cells/microwell) following the manufacturers protocol. EBs were incubated at 37°C with 5% CO2, with minimal disruption during the first 48 hours. Media changes, 50-75% of the total volume, were performed every 2 days. On Day 4, EBs were harvest according to the manufacturer protocol and transferred to a 10 cm tissue culture dish (Eppendorf, catalog 0030702018). On Day 5, Induction Medium was added to each well and incubated for 48 h at 37°C. On Day 7, high quality EBs (smooth and optically translucent edges) were embedded in Matrigel (Corning). EBs were transferred to a 15mL conical tube and resuspended in Expansion Medium. Using a P100 pipette, 67uL containing ∼20-30 EBs in medium were transferred to a microcentrifuge tube. Next, 100uL of thawed Matrigel was added to the microcentrifuge tube (3:2 ratio) and mix with the medium and EBs by pipetting up and down multiple times. Using a cut tip, the Matrigel-EB mixture was pipetted and spread onto the center of an ultra-low attachment six-well plate. Matrigel coat should fully envelop the EBs in 3D (thickness of >1mm). EBs should be evenly distributed to avoid contacting each other or the wall of the well. Incubate the Matrigel at 37°C for 30 min to solidify. Gently add 3mL of Expansion Medium and incubate at 37°C for 3 days.

On Day 10, the Matrigel coat was broken by vigorously pipetting up and down. Organoids were transferred to a 15mL conical tube. Healthy organoids will precipitate faster than Matrigel clumps and organoid debris. By gently mixing, debris can be removed with supernatant after the healthy organoids precipitate to the bottom of the tube. Organoids were then resuspended in Maturation media and distribute into an ultra-low attachment 12-well plate (6-10 organoids per well). Maturation media was added to a final volume of 3mL/well. Subsequently, the plate was inserted into the bottom frame of the Spin∞and the lid was positioned in place. The whole system was ten moved to a 37°C incubator and the shaking speed was set at 90rpm. Full Maturation Medium change was performed every 3–4 days. Reconstitution of extracellular matrix was performed on Day 40. Matrigel was thawed on ice and dissolved in Maturation Media in a 1:50 dilution. Transmitted-light images were acquired using an EVOS® XL Core Imaging System. The software used for processing was ImageJ.

### Tissue preparation and Immunohistochemistry

Cerebral organoids were fixed in 4% Paraformaldehyde in Phosphate Buffered Saline (PBS) for 15-20 min at 4°C. Organoids were washed 3 times with PBS and then incubated in 30% sucrose solution overnight at 4°C. Organoids were embedded in 7.5% gelatin/10% sucrose solution (Sigma, catalog G1890-100G and S7903-250G) and sectioned with a cryostat (Leica CM1950) at 15um thickness.

For immunostaining, freezing medium was washed with PBS before permeabilization with 0.2% Triton-X in PBS for 1 hr. Tissues were blocked with blocking medium consisting of 10% donkey serum in PBS with 0.1% Tween-20 (PBST) for 30 min. Primary antibodies diluted in blocking solution were applied to the sections overnight at 4°C. After washing 3 times with PBST, secondary antibodies were diluted in blocking solution were applied to the sections for 1 hr at room temperature. Finally, sections were washed 3 times with PBST and stained with Hoechst. Confocal images were acquired using an Andor DU-897 EMCCD camera mounted on a Nikon Spinning Disk Microscope. The software used for image acquisition and reconstruction was NIS-Elements Viewer (Nikon).

### Building the Spin∞ bioreactor

The procedure for assembling and operating the Spin∞ bioreactor is described below and in Figures 1-16. Figure 1 shows all components. Additional details on hardware and all components of the Spin∞ are provided in Supplemental Information.

#### Hardware assembly and operation

##### Motor preparation

1. Solder positive and negative terminals (3 ft of 2 wire cable).
2. Heat shrink tubing to each terminal.
3. Crimp on male version of the JST plugs.
4. Add 6 screws loosely to motor (Figure 2).
5. Optional: coat the entire motor unit with parylene.

##### Acrylic plate preparation

1. Laser cut the acrylic plate with appropriate dimensions (Figure 3). Settings will vary based on laser wattage. Here, a 45W laser cutter was used with 80% power and 70% speed with two passes.

##### Bioreactor assembly

1. Optional: coat the 3D printed 12-Well Plate Lid and Base with parylene.
2. Optional: attach the paddles and the gears to the 3D printed holders (Parylene Template for Gears and Parylene Template for Paddles) and coat with parylene C. Be sure to place the bottom of the gears facing up to maximize vapor deposition.
3. Insert the stainless steel M3 nuts in the gears.
4. Use a 5 mm (diameter of PTFE collar) drill bit to gently core out the center of the PTFE collar and insert into the 12-Well Plate Lid. Be sure to lightly drill out the center of the PTFE collar and remove as little as possible.
5. Insert CW Paddles and CCW Paddles into the lid and PTFE collars (Figure 4). Proper position of each paddle is noted.
6. To attach the Gears, start with the Motor Shaft Gear at the B2 position (Figure 5).

a. The bottom set screw gear needs to be aligned with the opening on the paddle.
b. Screw in the top screw. Do not screw the screws in any further than what is shown in the image. The shaft should easily and freely spin (Figure 6).
7. Attach the other gears to the rest of the wells working counterclockwise around the lid (Figure 5). The teeth of the gears should be aligned and interlock.
8. Attach the hex standoffs and 4 hex nuts to the base (Figure 7).

a. Screw on the 45 mm hex standoffs with washers attached to the 4 locations with the hex nuts on the base.
b. Attach screws to hex standoffs.
9. Attach the hex standoffs to the lid and insert the nuts into the 12-Well Plate Lid (Figure 8). Screw the 35 mm hex standoffs with washers attached to all the locations with the hex nuts on the lid and attach screws to the hex standoffs.
10. Autoclave the 3D printed and stainless-steel parts either assembled or as individual pieces for 60 minutes (Figure 9).

a. Place all autoclavable items in an autoclavable bag upside down, and seal the bag.
b. Everything but the motor, acrylic plate, and cell culture plate can be autoclaved.
11. After autoclaving is complete, spray the autoclave bag with 70% ethanol and transfer it into the biosafety hood. Take the device out of the bag.
12. Outside of the hood, insert the shaft of the motor through the hole in the acrylic plate, and attach the motor to the acrylic plate using the six 10 mm screws (Figure 10).
13. Transfer the motor and acrylic plate into the biosafety hood. The motor and acrylic plate should be sprayed with 70% ethanol for sterilization.
14. Remove the lid from a sterile 12-well plate and place it into the Base. Add the 12-Well Plate Lid to the top of the plate (Figure 11).
15. Take the screws off the hex standoffs. The screws can be stored in a sterile petri dish.
16. Make sure that the top screw on the Motor Gear Shaft is loosened.
17. Place the acrylic sheet on top of the hex standoffs. Make sure that the motor shaft goes inside the Motor Shaft Gear.
18. Attach the motor to the Motor Gear Shaft (Figure 12).

a. Find the beveled side of the motor shaft and line it up with the top screw of the Motor Shaft Gear.
b. By hand, screw the top screw to where there is a little play with the set screw in the shaft.
c. Critical: avoid overtightening the top screw of the Motor Gear Shaft.
19. Thread the screws into the 35 mm hex standoffs (12-Well Plate Lid) through the acrylic plate (Figure 13). Do not tighten completely! There should be some play.
20. Thread the screws through the acrylic plate to the 45 mm hex standoffs on the Base, and tighten all four screws. These should be securely tightened.
21. Tighten the screws attached to the 35 mm hex standoffs. Once fully tightened, turn the screws ½ counter clockwise. The screws to the 12-Well Plate Lid hex standoffs are designed to be loose.
22. Before using with cells, allow the assembled bioreactor to run for at least 24-48 hours dry in a 37°C incubator to remove any debris from the 3D printed parts. Sterile PBS can then be used to wash the bioreactor.

#### Electronics assembly

##### Initial assembly

1. Follow the Raspberry Pi Touchscreen assembly instructions with the Raspberry Pi 3 A+ and Raspberry Pi SD card preloaded with Noobs. The Raspberry Pi should be powered by the 5V microUSB power source provided in the kit.
2. Once the touchscreen has been assembled, the electronics need to be assembled as described below.
3. First, remove the jumper from enable A and B on the L298n bridge (Figure 14).
4. Connect the bridge to the GPIO pins on the Raspberry Pi (Figure 15). For the first motor, connect the ENA pin on the L298n bridge to GPIO pin 11, connect IN1 on the L298n bridge to GPIO pin 16 and connect the IN2 on the L298n bridge to GPIO pin 18 (Figure 15). Repeat this for all motors and L298n bridges. Each L298n bridge can control two motors, and using the instructions in Figure 15, a single Raspberry Pi can run up to five motors.
5. Connect the in-line power toggle to turn the bridge on and off.
6. Make sure the L298n bridge has a 12V input. The ground wire goes to the bridge and the Raspberry Pi. This is essential and will not work without both grounds.
7. Last, connect 3 feet of 2-wire to the output screw terminals for motor 1 and attach a female JST connector on the other end.

##### Software download and installation

1. Turn on the Raspberry Pi and open the command terminal.
2. Type the following text in the command line (Figure 16A-B): git clone https://github.com/LippmannLab/Spinfinity.git
3. Enter (Figure 16C): cd/home/pi/Spinfinity.
4. A new line should read (Figure 16D): pi@raspberrypi:∼/Spinfinity $
5. Enter (Figure 16E): python 5Motors.py
6. The program will now open (Figure 16F). If the Raspberry Pi is restarted, then open the command terminal again and re-enter steps 3-5 to open the program.

#### Procedure for media changes

1. Turn off the motor using the in-line power switch and disconnect the JST connector.
2. Take the Spin∞ out of the incubator and place it in a biosafety hood.
3. Unscrew the screws attached to the 45 mm hex standoffs. Do not lose the screws.
4. Transfer the 12-Well Plate Lid (that is still attached to the acrylic plate and motor) to a sterile, empty 12 well plate. This will expose the plate of organoids resting in the Base, which can be removed for imaging if desired.
5. Change the media:

a. Tilt the plate approximately 45 degrees so that the organoids will sink to the bottom of the well.
b. Carefully remove the media from each well using a P1000 micropipette and put the spent media into an empty 15 mL conical tube.
c. Check the spent media to make sure that no organoids were accidentally removed.
d. Once spent media is removed, add 3-4 ml of fresh media to each well using a P1000 micropipette.
6. Return the 12-Well Plate Lid on top of the 12-well plate that is resting in the Base.
7. Screw the screws back onto the 45 mm hex standoffs tightly.
8. Put the device back into the incubator.
9. Connect the JST connector and turn on the motor. Check to make sure that the gears and paddles are turning at the desired speed.

### Statistical analysis

Data was verified across all biological replicates, where a minimum of three independent experiments was performed. Statistical significance was determined by one-way ANOVA.

## Acknowledgments

We thank Dr. Bryan Millis (Vanderbilt Nikon Center for Excellence) for his technical support with image acquisition and processing, Dr. Jenny Schafer (Vanderbilt Cell Imaging Shared Resource) for support with light sheet imaging of embryoid bodies, and Dr. Rebecca Ihrie for helpful technical and scientific insight. We would like to thank members of the Gama and Lippmann Laboratories for helpful discussions and comments to the manuscript. Funding was provided by a Ben Barres Early Career Acceleration Award from the Chan Zuckerberg Initiative (to ESL); 1R35GM128915-01 (to VG); 1R21 CA227483-01A1 (to VG); the Precision Medicine and Mental Health Initiative sponsored by the Vanderbilt Brain Institute (to VG and ESL); and NSF 1506717 (to LB). BJO was supported by the Vanderbilt Interdisciplinary Training Program in Alzheimer’s Disease (T32 AG058524). KMB was supported by the Training Program in Environmental Toxicology (T32 ES007028). Image acquisition and analysis were performed in part through the use of the Nikon Center of Excellence within the Vanderbilt Cell Imaging Shared Resource (supported by NIH grants CA68485, DK20593, DK58404, DK59637 and EY08126), Vanderbilt University Medical Center’s Translational Pathology Shared Resource supported by NCI/NIH Cancer Center Support Grant 2P30 CA068485-14, and the Vanderbilt Mouse Metabolic Phenotyping Center Grant 5U24DK059637-13.

The authors have no conflict of interest to declare.

