## Supplemental Information for "Spin∞ an improved miniaturized spinning bioreactor for the generation of human cerebral organoids from pluripotent stem cells"

#### 1. Hardware description

The Spin∞ is built primarily from 3D printed ULTEM 1010 resin to permit autoclaving. Alternate 3D printing filaments, such as acrylonitrile butadiene styrene (ABS), can be used but require more extensive sterilization steps such as sequential washes in 10% bleach, 70% ethanol, distilled water washes and UV irradiation. The bottom 3D printed frame contains an inset to hold a 12-well plate, as well as inserts for the metal standoffs. The top 3D printed frame houses PTFE collars, gears, and paddles, which are manually assembled. All of the gears except one are used solely for turning, while the remaining gear connects directly to a DC 12V 100RPM gear motor (Greartisan) attached to an acrylic plate. The acrylic plate rests on the metal standoffs and is screwed into place to ensure mechanical stability. All components can be optionally coated with parylene to prevent corrosion of the metal housing on the motor and to add an additional hydrophobic barrier to prevent absorption of media into the 3D printed parts. The motor connects to a L298n bridge and a Raspberry Pi 3A+ that controls the motor speed through a touchscreen interface. **Figure 1** shows all components.

- Increased motor life span under high temperature and humidity conditions.
- Can be autoclaved after assembly, thus removing the need for disassembly for cleaning and sterilization.
- Sturdy, fully enclosed assembly to prevent leaks and spills while ensuring consistent operation.
- Integrated touch screen for changing motor speed.

### 2. Design files

**Table 1** contains links to all STL design files for 3D printed parts, the python script for running the bioreactor, and a video demonstration of step-by-step assembly of the device.

**Table 1. Design files.**

| Design file name | File type | Open source license | Location of the file |
|---|---|---|---|
| Base | STL | CC-BY-SA 4.0 | https://osf.io/tavck/?view_only=e89148170d1046bebec51977b3177180 |
| 12-Well Plate Lid | STL | CC-BY-SA 4.0 | https://osf.io/adqye/?view_only=e89148170d1046bebec51977b3177180 |
| CCW Paddle | STL | CC-BY-SA 4.0 | https://osf.io/qfxsy/?view_only=e89148170d1046bebec51977b3177180 |
| CW Paddle | STL | CC-BY-SA 4.0 | https://osf.io/8pshg/?view_only=e89148170d1046bebec51977b3177180 |
| Gear | STL | CC-BY-SA 4.0 | https://osf.io/abgmr/?view_only=e89148170d1046bebec51977b3177180 |
| Motor Shaft Gear | STL | CC-BY-SA 4.0 | https://osf.io/9dnbq/?view_only=e89148170d1046bebec51977b3177180 |
| Parylene Template for Gears *(Optional)* | STL | CC-BY-SA 4.0 | https://osf.io/wruvb/?view_only=e89148170d1046bebec51977b3177180 |
| Parylene Template for Paddles *(Optional)* | STL | CC-BY-SA 4.0 | https://osf.io/u89v3/?view_only=e89148170d1046bebec51977b3177180 |
| X4 L298n Bridge Holder *(Optional)* | STL | CC-BY-SA 4.0 | https://osf.io/9nyr2/?view_only=e89148170d1046bebec51977b3177180 |
| 5Motors | PY | CC-BY-SA 4.0 | https://osf.io/trca4/?view_only=e89148170d1046bebec51977b3177180 |
| Spinfinity Assembly | AVI | CC-BY-SA 4.0 | https://osf.io/kb3g9/?view_only=e89148170d1046bebec51977b3177180 |

- The **Base** holds a 12-well cell culture plate where the organoids are cultured.
- The **12-Well Plate Lid** holds the PTFE collars, CCW Paddles, CW Paddles, Gears, and Motor Shaft Gear. This component replaces the traditional lid on a cell culture plate.
- The **CCW Paddle** is partially submerged in the media and carries out mixing in a counter clockwise direction.

- The **CW Paddle** is partially submerged in the media and carries out mixing in a clockwise direction.
- The **Gear** holds the paddles and spins under the control of the motor.
- The **Motor Shaft Gear** directly attaches to the motor and turns all of the other gears.
- The **Parylene Template for Gears** holds the gears during parylene deposition. This print is optional if the user will not coat with parylene.
- The **Parylene Template for Paddles** holds the paddles during parylene deposition. This print is optional if the user will not coat with parylene.
- The **X4 L298n Bridge Holder** holds four individual bridges for driving the motors.
- The **5Motors** is a python script to enable touchscreen control over motor speed and direction.
- The **Spinfinity Assembly** is a video showing each step of the bioreactor installation.
- All files can be found under the reserved Spinfinity repository: doi:10.17632/89g9nsd4gk.1

**3. Bill of Materials**

**Tables 2-4** contain descriptions, suppliers, and pricing on hardware, 3D prints, and software components.

**Table 2. Hardware components.**

| Quantity | Part Name | Supplier | Part Number | Cost |
|---|---|---|---|---|
| 1 | Acrylic Sheet 12" X 12" X ¼" | McMaster-Carr | 8560K354 | $17.34 |
| 4/Bioreactor | 18-8 Stainless Steel 45 mm Hex | McMaster-Carr | 93655A226 | $4.25 |
| 4/Bioreactor | 18-8 Stainless Steel 35 mm Hex | McMaster-Carr | 93655A224 | $4.11 |
| 8/Bioreactor | 18-8 Stainless Steel Washer M3 | McMaster-Carr | 93475A210 | $1.62 (Pack of 100) |
| 8/Bioreactor | 18-8 Stainless Steel Hex Nut M3 0.5mm Thread | McMaster-Carr | 91828A211 | $5.55 (Pack of 100) |
| 14/Bioreactor | 18-8 Stainless Steel Philips Flat Head Screw M3 | McMaster-Carr | 92010A120 | $4.65 (Pack of 100) |
| 12/Bioreactor | PTFE Collars | McMaster-Carr | 2685t11 | $6.16 |
| 1/Bioreactor | 12-Well Cell Culture Plate | Corning | 3737 | $431.00 (Case of 100) |
| 1/Bioreactor | Autoclavable Bags | Fisher Scientific | 01-812-58 | $182.00 (Pack of 100) |
| 1/Bioreactor | Motor 100RPM, 12V, Eccentric Shaft | Amazon | B0721T1PXQ | $15.49 |
| 1 Set | Screwdriver Set | McMaster-Carr | 52985A22 | $30.62 |
| 1 Set | Solder | McMaster-Carr | 7667A51 | $38.77 |
| 1 Set | Heat Shrink Tubing | McMaster-Carr | 6334K414 | $12.71 |

**Table 3. 3D print components.**

| Quantity | Part Name | Supplier |
|---|---|---|
| 1 | 12-Well Plate Lid | Xometry (ULTEM 1010) |
| 1 | Base | Xometry (ULTEM 1010) |
| 1 | Motor Shaft Gear | Stratasys (ULTEM 1010) |
| 11 | Gear | Stratasys (ULTEM 1010) |
| 6 | CW Paddle | Stratasys (ULTEM 1010) |
| 6 | CCW Paddle | Stratasys (ULTEM 1010) |
| 1 | Parylene Template for Gears | Stratasys (ABS) |
| 1 | Parylene Template for Paddles | Stratasys (ABS) |
| 1 | L298n Bridge Holder | Stratasys (ABS) |

- ULTEM 1010 can be autoclaved and is recommended for 3D prints that will be used under sterile conditions. ABS can be used instead, but several additional steps are required to sterilize the 3D printed parts (parylene coating and a series of 10% bleach, 70% ethanol, distilled water washes, as well as UV radiation).
- The 3D prints for Parylene Template for Plate Gears, Parylene Template for Plate Paddles, and L298n Bridge Holder can be made from ABS because they are not used under sterile conditions.
- The Parylene Template for Plate Gears and Parylene Template for Plate Paddles do not need to be printed if the user does not intend to coat with parylene.
- Cost of 3D printed parts depends on quantity ordered and vendor.

**Table 4. Electronic components.**

| Quantity | Part Name | Supplier | Part Number | Unit Cost |
|---|---|---|---|---|
| 1/Bioreactor | Raspberry Pi 3 A+ | Sparkfun | DEV-15139 | $29.95 |
| 1/Bioreactor | Raspberry Pi 3 A+ Power supply | Sparkfun | TOL-13831 | $7.95 |
| 1/Raspberry Pi | Raspberry Pi™ - 16GB MicroSD NOOBS Card | Sparkfun | COM-13945 | $24.95 |
| 1/Raspberry Pi | Raspberry Pi LCD - 7" Touchscreen | Sparkfun | LCD-13733 | $64.95 |
| 1/Bioreactor | Power Supply 12V, 5A | Amazon | B06Y64QLBM | $8.99 |
| 1/8 Bioreactors | 4-way DC Jack Splitter | Amazon | B00MHUGL7W | $8.29 |
| 1/Bioreactor | 5 PCS L298N Motor Drive Controller Board | Amazon | B06X9D1PR9 | $15.99 (Pack of 5) |
| 1/Bioreactor | 100ft 4 Pin RGB Extension Cable Wire | Amazon | B074H7DM4B | $16.99 |
| 1/Motor | 2-pin JST SM Male & Female Plug Housing | Amazon | B0188DMF3A | $14.99 |
| 1/L298N Bridge | In-line Power Toggle (*Optional)* | Amazon | B0782JXQNP | $5.85 |
| 1/LCD Screen | SmartPi Touch Case *(Optional)* | Amazon | B01HV97F64 | $27.99 |
| 1/LCD Screen | Wireless Bluetooth Keyboard | Amazon | B00BX0YKX4 | $24.99 |

### 4. Build Instructions

The essential bioreactor components are 3D printed. The remaining hardware and electronics can be purchased from common suppliers.

The additional tools required for assembly include:

- Drill
- 5 mm drill bit
- Philips screwdriver
- $CO_2$ laser cutter
- Optional: parylene vapor deposition machine such as a Labcoter supplied by Specialty Coating Systems
- Solder and soldering iron
- Heat shrink tubing and heat shrink gun
- Crimper
- JST connectors (male and female)
- 2-wire (18-20 AWG, 66ft)
